# TOPBP1-Dependent DNA Damage Processing Promotes Centromere Loss in an Inviable *Xenopus* Hybrid

**DOI:** 10.64898/2026.09.16.752195

**Authors:** Cordell Clark, Riki Terui, Gheorghe Chistol, Rebecca Heald

## Abstract

Hybrid incompatibility is associated with chromosome instability, DNA damage, and embryonic lethality, yet how cells respond to genome instability in hybrids is unclear. In inviable hybrids generated by fertilizing Xenopus tropicalis eggs with Xenopus laevis sperm, CENP-A is lost from paternal chromosomes, leading to elimination of two chromosome arms. Using hybrid Xenopus egg extract reactions, we show that CENP-A is removed from X. laevis chromosomes exposed to X. tropicalis cytoplasm through an RNA polymerase II-dependent process associated with DNA damage. Topoisomerase II Binding Protein 1 (TOPBP1) localized to damaged acentric chromosomes during metaphase, and its depletion impaired chromosome alignment, increased DNA damage, and blocked CENP-A removal. Inhibition of DNA polymerase theta increased DNA damage while promoting CENP-A retention, indicating that DNA damage processing rather than damage itself promotes CENP-A loss. Together, our findings reveal separable roles for TOPBP1 in chromosome organization and centromere destabilization, showing how a protective genome surveillance pathway can instead drive chromosome instability in an inviable hybrid

## Introduction

Embryonic lethality is a common barrier to successful hybridization between closely related species, contributing to speciation and reproductive isolation across eukaryotes. In many hybrids, incompatibility is accompanied by genome instability, including DNA damage and chromosome instability. Studies in diverse experimental systems have implicated whole genome elimination (Fujiwara et al., 1997), defects in chromatin packaging of repetitive elements (Ferree and Barbash, 2009; Jagannathan and Yamashita, 2021), and the inability to resolve topological DNA stress (Brand and Levine, 2022; Brand et al., 2026). These findings suggest that chromosomes can become unstable when exposed to a divergent cellular environment, but whether such defects engage common molecular response pathways remains unclear.

Observations across mammalian, insect, plant, and *Xenopus* models point to centromere dysfunction as a recurring source of genome instability in inviable hybrids (El Yakoubi and Akera, 2023; Gibeaux et al., 2018; Kitaoka et al., 2022; Maheshwari et al., 2015; Rosin and Mellone, 2016; Sanei et al., 2011). Central to centromere maintenance is CENP-A, the histone H3 variant that epigenetically defines centromeres within chromosome regions often enriched in repetitive DNA (McKinley and Cheeseman, 2016). Along with members of the constitutive centromere-associated network, CENP-A coordinates kinetochore assembly, mediating the physical interaction between mitotic chromosomes and spindle microtubules necessary for accurate chromosome segregation (Black et al., 2007; Foltz et al., 2006; Hara et al., 2023; Kato et al., 2013; Milks et al., 2009). CENP-A chromatin may also protect centromere integrity by buffering against replication stress and limiting DNA damage (Giunta and Funabiki, 2017; Giunta et al., 2021). Paradoxically, despite their essential functions in genome maintenance, centromeres are rapidly evolving regions that are particularly susceptible to DNA damage (Altemose et al., 2022; Logsdon et al., 2024; Henikoff et al., 2001; Malik and Henikoff, 2001; Saayman et al., 2023; Showman et al., 2024). Thus, defining how centromere function is disrupted in hybrids could reveal fundamental mechanisms that preserve CENP-A stability and how evolutionary divergence can compromise them.

Our group has previously used the *Xenopus* egg extract system to show that a subset of *Xenopus laevis* sperm chromosomes lose CENP-A when replicated and cycled into metaphase in *Xenopus tropicalis* extract (Gibeaux et al., 2018; Kitaoka et al., 2022). Similar to observations in *Drosophila*, this CENP-A defect could be rescued by addition of *X. laevis* CENP-A and its dedicated chaperone, HJURP (Kitaoka et al., 2022; Rosin and Mellone, 2016). Because CENP-A deposition is tightly regulated and essential for kinetochore assembly in vertebrates (Conti et al., 2024; Flores Servin et al., 2023; Jansen et al., 2007; McKinley and Cheeseman, 2014; Moree et al., 2011), these findings implicated defective CENP-A maintenance in hybrid centromere instability, but did not distinguish between impaired deposition and active CENP-A removal. Moreover, why only a subset of *X. laevis* chromosomes is affected remains unclear.

Despite consistent CENP-A loss *in vitro* and loss of whole chromosome arms in vivo, *X. tropicalis* egg × *X. laevis* sperm hybrid embryos undergo normal cleavage divisions before dying at gastrulation and maintain a low, stable level of micronuclei during embryogenesis (Gibeaux et al., 2018; Kitaoka et al., 2022). This is surprising given that prior to the mid-blastula transition, embryonic cells lack gap phases and a functional spindle assembly checkpoint (Hara et al., 1980; Vázquez-Diez et al., 2019; Zhang et al., 2015). Thus, acentric chromosomes should be mis-segregated each cell cycle, leading to progressive accumulation of micronuclei during development. How rapidly dividing hybrid embryonic cells limit micronucleus formation despite ongoing chromosome instability remains unknown.

One factor that could recognize and promote the segregation of acentric chromosomes in hybrids is Topoisomerase II binding protein 1 (TOPBP1), a multifunctional scaffold protein with essential roles in DNA replication initiation, DNA damage signaling, and DNA repair (Broderick et al., 2015; Day et al., 2024; De Marco Zompit et al., 2022; Leimbacher et al., 2019; Terui et al., 2024). Recent studies in human cells have shown that TOPBP1 tethers acentric chromosome fragments during mitosis, promoting their alignment and segregation while limiting micronucleus formation (Adam et al., 2021; Lin et al., 2023; Trivedi et al., 2023). In addition, TOPBP1 promotes the repair of DNA damage through multiple pathways, including break-induced replication and microhomology-mediated end joining (MMEJ) involving DNA Polymerase theta (POLθ) (Gelot et al., 2023; Martin et al., 2025). Whether these functions of TOPBP1operate during early embryogenesis or influence chromosome instability in hybrids are unexplored questions.

In this work, we leverage hybrid *Xenopus* egg extract reactions to investigate the mechanisms and consequences of CENP-A loss in inviable *X. tropicalis* × *X. laevis* hybrids. We show that CENP-A loss from a subset of *X. laevis* chromosomes is not due to defective assembly, but instead results from an active, transcription-dependent process in *X. tropicalis* cytoplasm. Our findings identify TOPBP1 as a key mediator of the response to transcription-induced DNA damage, with distinct roles in chromosome alignment and POLθ-dependent CENP-A removal. Together, our findings reveal how DNA damage processing can drive selective centromere destabilization in an inviable hybrid.

## Results

### CENP-A is actively removed from a subset of *Xenopus laevis* chromosomes

Our group previously showed that a subset of *X. laevis* sperm chromosomes loses CENP-A after replication and entry into metaphase in *X. tropicalis* extract (Gibeaux et al., 2018; Kitaoka et al., 2022). To distinguish between failed CENP-A deposition and active removal, we took advantage of the fact that CENP-A deposition occurs during early interphase in *Xenopus* egg extracts (Bernad et al., 2011; Flores Servin et al., 2023; Brown et al., 2026). *X. laevis* sperm nuclei were replicated in interphase-arrested *X. laevis* extract to allow normal CENP-A deposition, then driven into metaphase by addition of metaphase-arrested *X. laevis* or *X. tropicalis* extract. Mitotic chromosomes were subsequently isolated and analyzed for CENP-A by immunofluorescence (Figure 1A). If CENP-A loss observed previously reflected defective deposition, chromosomes that acquired CENP-A during interphase in *X. laevis* extract should retain the CENP-A upon subsequent exposure to *X. tropicalis* extract. Instead, a subset of chromosomes lost CENP-A following addition of metaphase *X. tropicalis* extract (Figure 1B,C), demonstrating that CENP-A is actively removed following exposure to *X. tropicalis* cytoplasm. Likewise, interphase nuclei isolated from *X. laevis* embryos in which CENP-A deposition had already occurred (Figure 1D) gave rise to a subset of mitotic chromosomes that lost CENP-A after exposure to *X. tropicalis* metaphase extract (Figure 1E,F). The frequency of CENP-A-negative chromosomes was comparable to that observed when chromosomes were cycled entirely in *X. tropicalis* extract (Gibeaux et al., 2018; Kitaoka et al., 2022), further demonstrating that CENP-A loss results from active removal rather than defective deposition.

**Figure 1.**
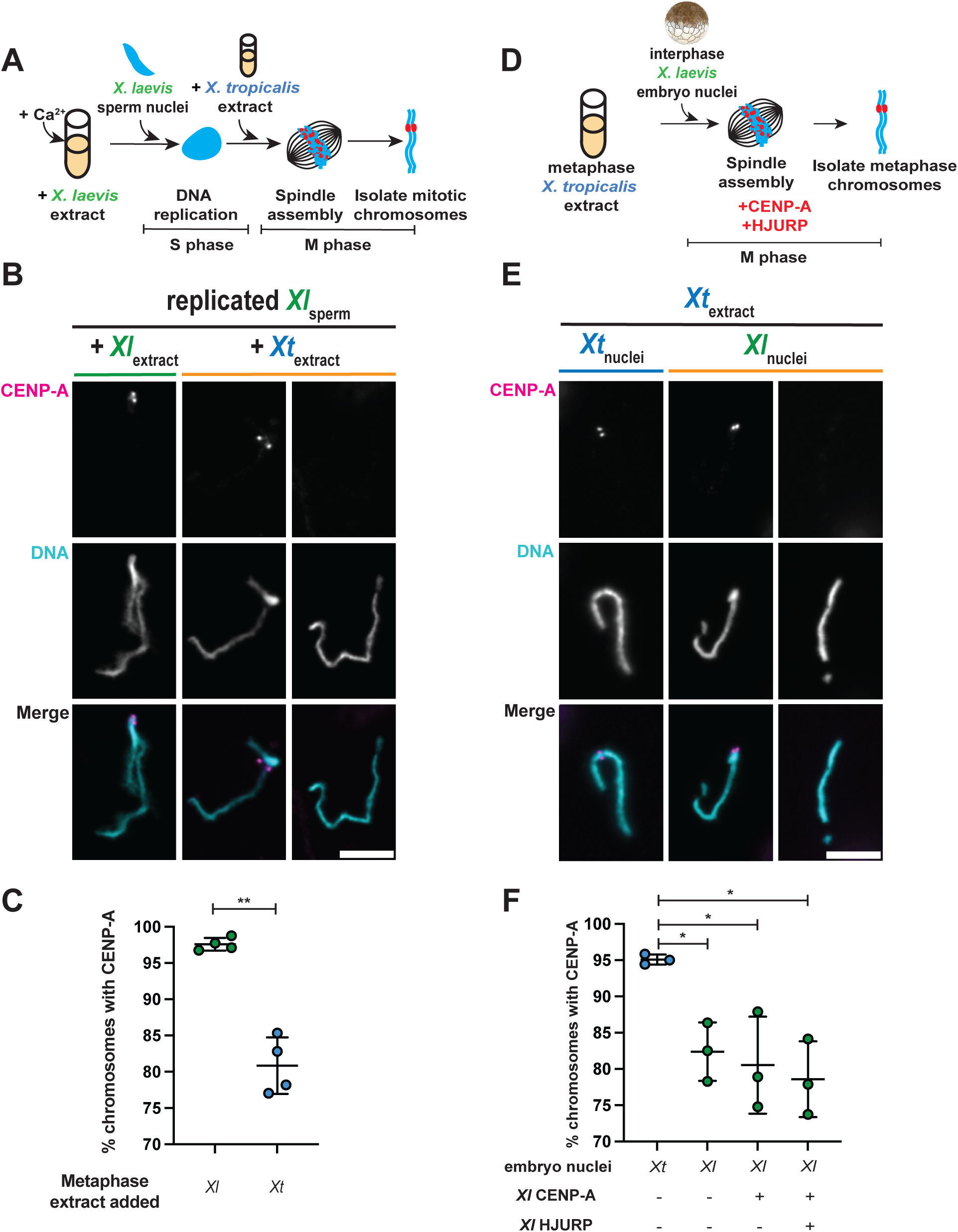
CENP-A is actively removed from a subset of *Xenopus laevis* chromosomes. (A) Schematic of the hybrid extract reaction using sperm nuclei. *X. laevis* sperm nuclei were replicated in interphase *X.* laevis *egg* extract and subsequently cycled into metaphase by addition of either *X. laevis* or *X. tropicalis* metaphase-arrested egg extract. Mitotic chromosomes were isolated and analyzed for CENP-A by immunofluorescence. (B) Representative images of replicated *X. laevis* sperm chromosomes cycled into metaphase in *X. laevis* or *X. tropicalis* extract. CENP-A, magenta; DNA, cyan. Scale bar, 5 µm. (C) Percentage of replicated *X. laevis* sperm chromosomes positive for CENP-A after cycling into metaphase in *X. laevis* or *X. tropicalis* extract. *n* = 3 independent extracts; >117 chromosomes analyzed per extract. P = 0.0025, Welch’s t test. (D) Schematic of the hybrid extract reaction using stage 8 embryo nuclei. Interphase nuclei isolated from stage 8 *Xenopus* embryos were cycled into metaphase in *X. tropicalis* egg extract. Mitotic chromosomes were isolated and analyzed for CENP-A by immunofluorescence. (E) Representative images of *X. tropicalis* or *X. laevis* chromosomes from stage 8 embryo nuclei cycled into metaphase in *X. tropicalis* extract. CENP-A, magenta; DNA, cyan. Scale bar, 5 µm. (F) Percentage of *X. tropicalis* or *X. laevis* chromosomes from stage 8 embryo nuclei positive for CENP-A after cycling into metaphase in *X. tropicalis* extract supplemented with the indicated centromere assembly factors. *n* = 3 independent extracts; >243 chromosomes analyzed per extract. P = 0.0456, 0.0229, and 0.0117, one-way ANOVA followed by Tukey’s multiple-comparisons test. Line indicates the mean, and the error bars indicate one standard deviation above and below the mean.

We next tested whether enhancing CENP-A chromatin assembly could prevent or reverse its loss. Addition of CENP-A and its chaperone HJURP during metaphase failed to restore CENP-A on embryo-derived chromosomes (Figure 1E,F), indicating that CENP-A cannot be reloaded once chromosomes have entered metaphase. In contrast, the addition of these factors during interphase prevented subsequent CENP-A loss when *X. laevis* sperm nuclei that were replicated in *X. laevis* extract were challenged with *X. tropicalis* metaphase extract (Figure S1A,B), consistent with previous results (Kitaoka et al., 2022). Notably, CENP-A was also lost from *X. laevis* nuclei incubated in interphase *X. tropicalis* extract (Kitaoka et al., 2022), and this loss could not be rescued by subsequent addition of *X. laevis* metaphase extract (Figure S1C,D). Together, these results demonstrate that CENP-A can be actively removed from a subset of *X. laevis* chromosomes following exposure to *X. tropicalis* cytoplasm during either interphase or metaphase. CENP-A loss can be prevented by enhancing its assembly during interphase but cannot be reversed after mitotic entry, consistent with evidence that CENP-A loading is inhibited in metaphase in *Xenopus* egg extracts (Flores Servin et al., 2023; Brown et al., 2026). Thus, centromere instability in these hybrid reactions results not from defective CENP-A assembly, but from selective, active removal of CENP-A from a subset of *X. laevis* chromosomes.

### CENP-A removal requires RNA Polymerase II activity

Having established that CENP-A is actively removed from a subset of *X. laevis* chromosomes upon exposure to *X. tropicalis* extract, we next asked whether this process requires RNA Polymerase II (RNAPII) activity. RNAPII-dependent centromeric transcription regulates CENP-A dynamics, supporting centromere maintenance under normal conditions while contributing to centromere destabilization when aberrantly regulated (Bouzinba-Segard et al., 2006; Chan et al., 2012; Chabot et al., 2024; Grenfell et al., 2016; McNulty et al., 2017). Consistent with previous findings (Kitaoka et al., 2022), continuous treatment with the RNAPII inhibitor triptolide throughout interphase and after transfer to *X. tropicalis* metaphase extract produced only a modest, statistically insignificant increase in CENP-A retention (Figure 2A–C). Because RNAPII activity contributes to CENP-A incorporation (Bobkov et al., 2018; Chabot et al., 2024; Rošić et al., 2014), continuous inhibition could mask an effect on subsequent CENP-A removal. We therefore restricted triptolide treatment to metaphase by adding the inhibitor to the *X. tropicalis* extract used to drive *X. laevis* sperm nuclei into metaphase following DNA replication and CENP-A deposition in *X. laevis* interphasic extract (Figure 2A). Under these conditions, RNAPII inhibition significantly increased CENP-A retention on mitotic chromosomes (Figure 2B,C). RNAPII inhibition similarly increased CENP-A retention in experiments using *X. laevis* embryo nuclei, both on mitotic chromosomes (Figure 2D–F), and in interphase nuclei (Figure S2A–D). Consistent with these results, inhibition of RNAPII with α-amanitin also increased CENP-A retention on embryo chromosomes (Figure S2E,F). Together, these findings demonstrate that CENP-A removal from a subset of *X. laevis* chromosomes in *X. tropicalis* cytoplasm requires RNAPII activity.

**Figure 2.**
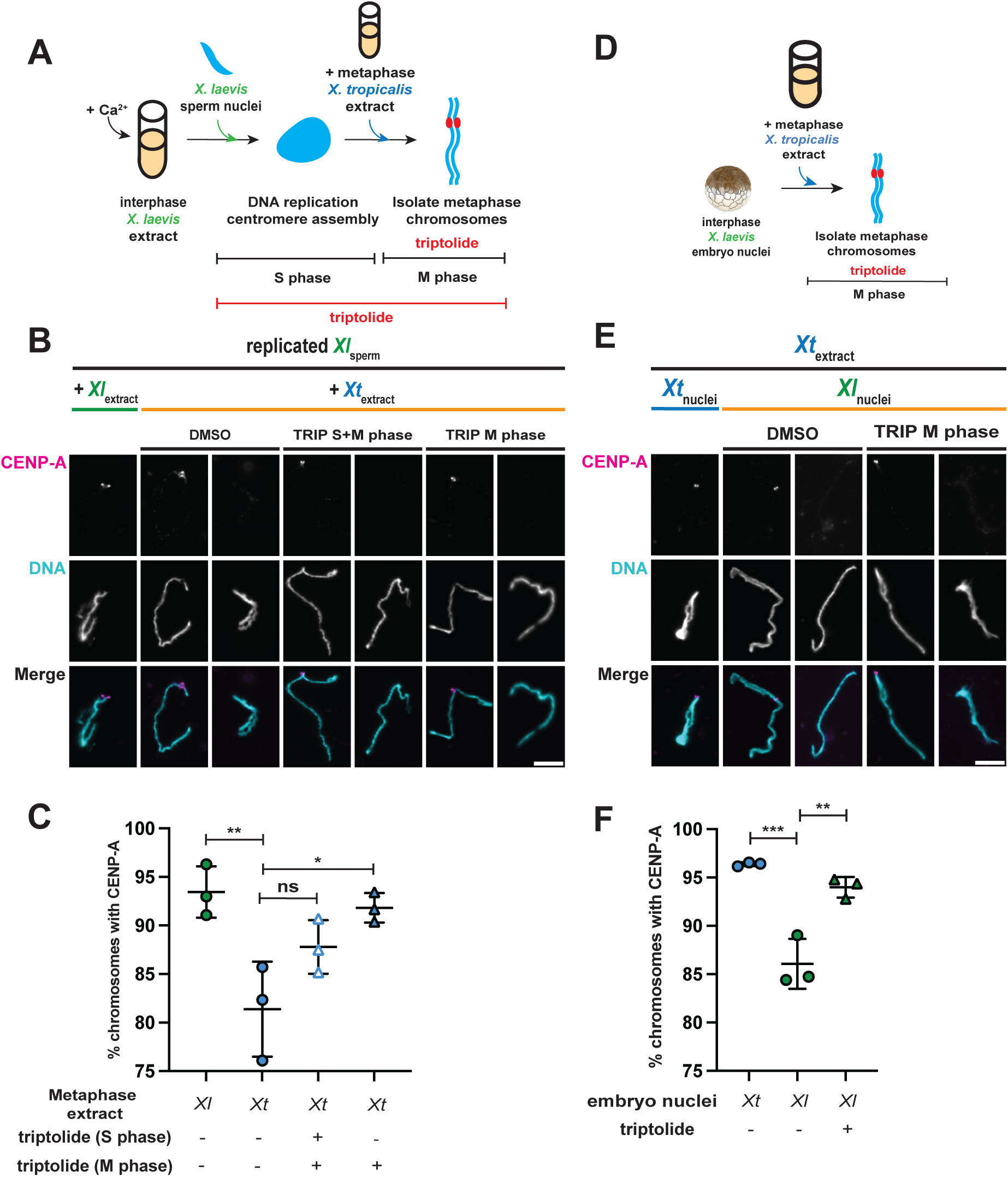
CENP-A removal requires RNA Polymerase II activity. (A) Schematic of hybrid extract reactions in which RNA polymerase II was inhibited with triptolide during both interphase and metaphase or during metaphase only. (B) Representative images of replicated *X. laevis* sperm chromosomes cycled into metaphase in *X. laevis* extract or *X. tropicalis* extract treated with DMSO, triptolide during both interphase and metaphase, or triptolide during metaphase only. CENP-A,magenta; DNA, cyan. Scale bar, 5 µm. (C) Percentage of replicated *X. laevis* sperm chromosomes positive for CENP-A after cycling into metaphase in *X. laevis* or *X. tropicalis* extract following treatment with DMSO (circles), triptolide throughout interphase and metaphase (white triangles), or triptolide during metaphase only (blue triangles). *n* = 3 independent extracts; >223 chromosomes analyzed per extract. P = 0.0025, 0.1671, and 0.0249, one-way ANOVA followed by Tukey’s multiple-comparisons test. (D) Schematic of hybrid extract reactions using stage 8 embryo nuclei treated with triptolide during metaphase. (E) Representative images of chromosomes from *X. tropicalis* embryo nuclei cycled into metaphase in species-matched *X. tropicalis* extract or *X. laevis* embryo nuclei cycled into metaphase in *X. tropicalis* extract treated with DMSO or 25 µM triptolide. CENP-A, magenta; DNA, cyan. Scale bar, 5 µm. (F) Percentage of chromosomes positive for CENP-A after *X. tropicalis* embryo nuclei were cycled into metaphase in species-matched *X. tropicalis* extract (blue circles) or *X. laevis* embryo nuclei were cycled into metaphase in *X. tropicalis* extract treated with DMSO (green circles) or 25 µM triptolide (green triangles). *n* = 3 independent extracts; >165 chromosomes analyzed per extract. P = 0.001 and 0.0018, one-way ANOVA followed by Tukey’s multiple-comparisons test. Line indicates the mean, and the error bars indicate one standard deviation above and below the mean.

### TOPBP1 localizes to damaged, acentric *X. laevis* chromosomes

Despite consistent CENP-A loss from a subset of *X. laevis* chromosomes in hybrid extract reactions, micronuclei remained relatively rare during early embryogenesis in *X. tropicalis* egg × *X. laevis* sperm hybrids (Gibeaux et al., 2018). We therefore asked whether chromosomes that had lost CENP-A were recognized by TOPBP1, a factor that promotes tethering of acentric chromosome fragments during mitosis in human cells (Lin et al., 2023; Trivedi et al., 2023). In species-matched reactions using *X. tropicalis* extract to drive *X. tropicalis* embryo nuclei into metaphase, TOPBP1 localized predominantly to spindle poles, with occasional puncta along the metaphase plate, consistent with previous observations in vertebrate cells (Bang et al., 2011; Pedersen et al., 2015; Reini et al., 2004) (Figure 3A). In contrast, spindles isolated from hybrid reactions using *X. tropicalis* extract to drive interphasic *X. laevis* embryo nuclei into metaphase exhibited prominent TOPBP1 staining on metaphase chromosomes and throughout the spindle (Figure 3A,B, S3A-D). Both the frequency of spindles containing TOPBP1-positive chromosomes and TOPBP1 fluorescence intensity at the metaphase plate were significantly reduced by RNAPII inhibition with triptolide (Figure 3A,B,S3E).

**Figure 3.**
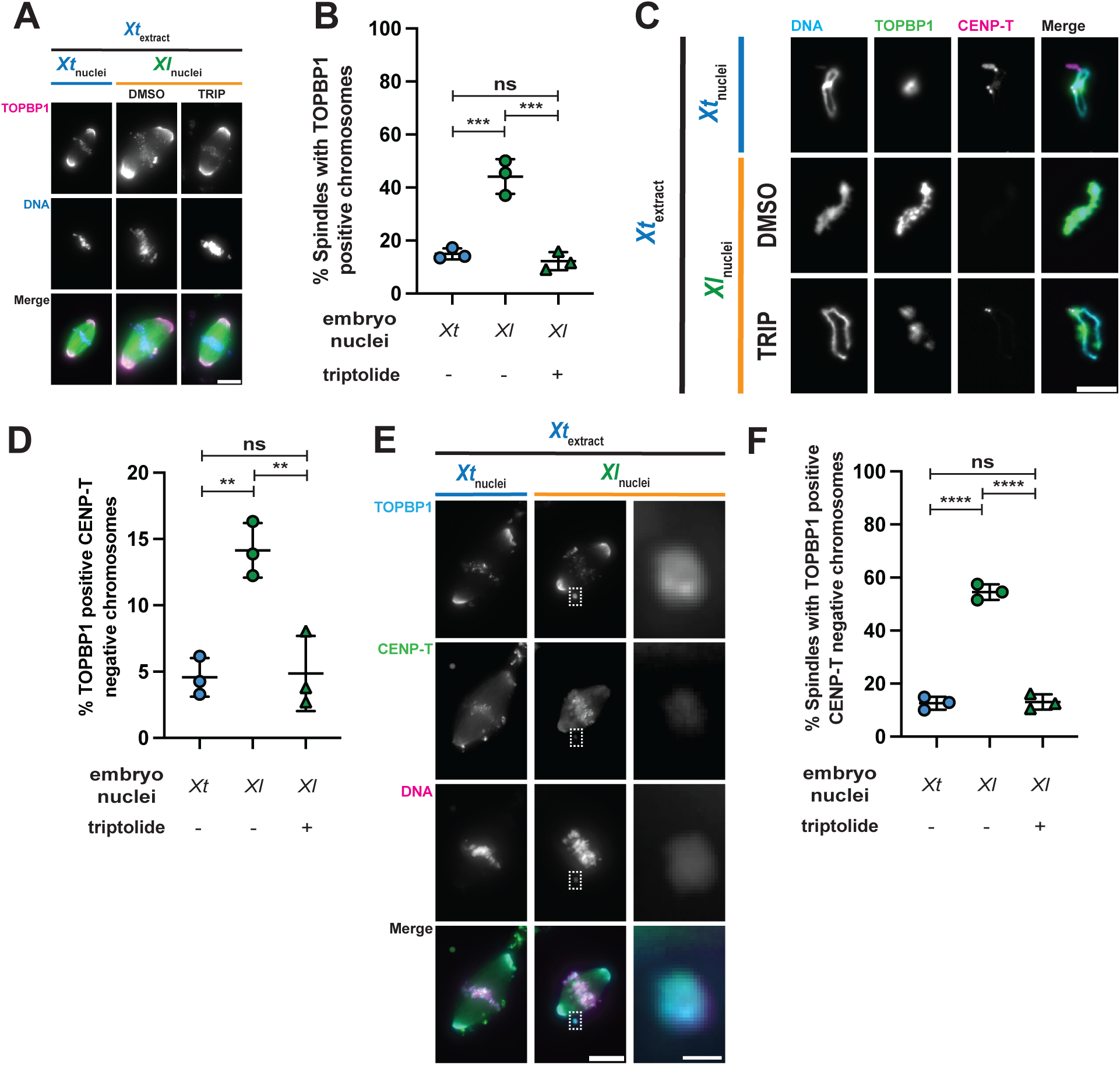
TOPBP1 localizes to acentric *X. laevis* chromosomes. (A) Representative immunofluorescence images of metaphase spindles assembled in *X. tropicalis* extract using *X. tropicalis* or *X. laevis* embryo nuclei treated with DMSO or 25 µM triptolide. TOPBP1, magenta; DNA, blue; tubulin, green. Scale bar, 10 µm. (B) Percentage of metaphase spindles displaying TOPBP1-positive structures along the metaphase plate in reactions using *X. tropicalis* embryo nuclei (blue circles) or *X. laevis* embryo nuclei treated with DMSO (green circles) or 25 µM triptolide (green triangles). *n* = 3 independent extracts; >140 spindles analyzed per extract. P = 0.0005, 0.0003, and 0.7467, one-way ANOVA followed by Tukey’s multiple-comparisons test. (C) Representative immunofluorescence images of mitotic chromosomes from *X. tropicalis* extract reactions using *X. tropicalis* embryo nuclei or *X. laevis* embryo nuclei treated with DMSO or 25 µM triptolide. DNA, cyan; TOPBP1, green; CENP-T, magenta. Scale bar, 5 µm. (D) Percentage of mitotic chromosomes displaying TOPBP1 staining along the chromosome axis and lacking CENP-T in reactions using *X. tropicalis* embryo nuclei (blue circles) or *X. laevis* embryo nuclei treated with DMSO (green circles) or 25 µM triptolide (green triangles). *n* = 3 independent extracts; >176 chromosomes analyzed per extract. P = 0.0042, 0.0048, and 0.9860, one-way ANOVA followed by Tukey’s multiple-comparisons test. (E) Representative immunofluorescence images of metaphase spindles assembled in *X. tropicalis* extract using *X. tropicalis* or *X. laevis* embryo nuclei, showing TOPBP1 and CENP-T localization. TOPBP1, cyan; CENP-T, green; DNA, magenta. Scale bar, 10 µm. Dashed box indicates the region enlarged at right, showing a TOPBP1-positive, CENP-T–negative chromosome. Scale bar, 1 µm. (F) Percentage of metaphase spindles displaying TOPBP1-positive, CENP-T–negative chromosome structures in reactions using *X. tropicalis* embryo nuclei (blue circles) or *X. laevis* embryo nuclei treated with DMSO (green circles) or 25 µM triptolide (green triangles). *n* = 3 independent extracts; >85 spindles analyzed per extract. P < 0.0001, P < 0.0001, and P = 0.9756, one-way ANOVA followed by Tukey’s multiple-comparisons test. Line indicates the mean, and the error bars indicate one standard deviation above and below the mean.

To determine whether these TOPBP1-positive chromosomes were acentric, we co-stained metaphase spindles and isolated chromosomes for TOPBP1 and the kinetochore protein CENP-T. TOPBP1-positive chromosomes frequently lacked CENP-T (Figure 3C–F), indicating that TOPBP1 associates with chromosomes lacking functional centromeres in hybrid extract reactions. TOPBP1 localization in species-matched reactions was largely restricted to chromosome ends and discrete puncta, whereas in hybrid reactions it extended along the length of a subset of chromosomes, with a frequency comparable to that of CENP-A loss. Triptolide treatment reduced the frequency of this chromosome-wide TOPBP1 staining (Figure 3C,D, S3A,B). Thus, TOPBP1 recruitment to acentric chromosomes in hybrid reactions is increased by RNAPII activity.

Because TOPBP1 also functions in response to DNA damage in mitosis (De Marco Zompit et al., 2022; Lin et al., 2023; Trivedi et al., 2023), we examined the DNA damage marker γH2AX to determine whether TOPBP1-positive acentric chromosomes were also associated with DNA damage. TOPBP1-positive metaphase chromosomes frequently co-stained for γH2AX in spindles assembled from *X. laevis* embryo nuclei in *X. tropicalis* extract (Figure 4A,B), indicating that TOPBP1 recognizes damaged, acentric chromosomes in hybrid reactions.

**Figure 4.**
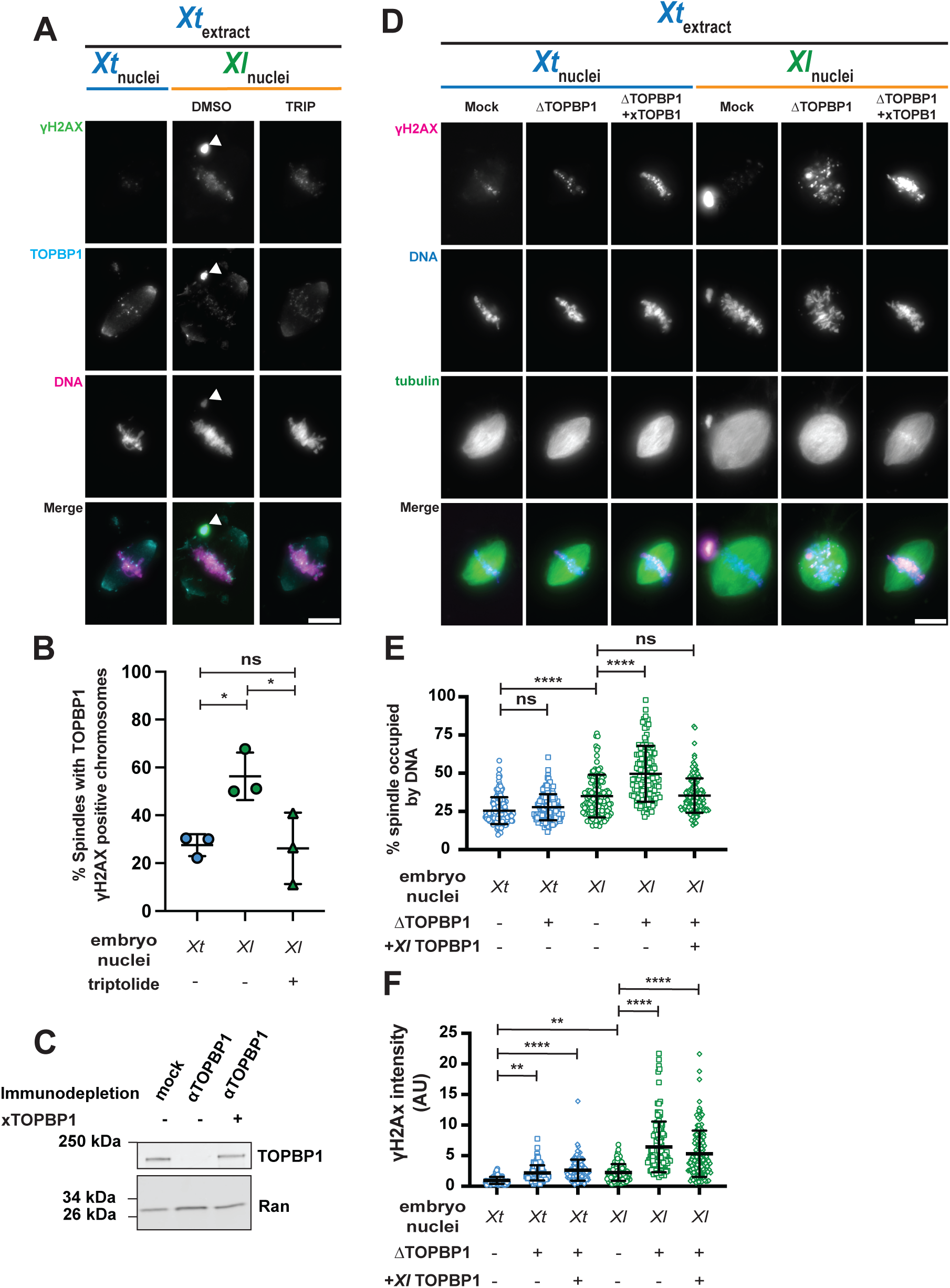
TOPBP1 promotes chromosome alignment and DNA damage processing in hybrid extract reactions. **(A**) Representative immunofluorescence images of metaphase spindles assembled in *X. tropicalis* extract using *X. tropicalis* embryo nuclei or *X. laevis* embryo nuclei treated with DMSO or 25 µM triptolide. Arrow indicates a chromosome with displaying TOPBP1 and γH2AX localization. γH2AX, green; TOPBP1, cyan; DNA, magenta; tubulin, grayscale. Scale bar, 10 µm. (B) Percentage of metaphase spindles displaying TOPBP1- and γH2AX-positive chromosomes in reactions using *X. tropicalis* embryo nuclei (blue circles) or *X. laevis* embryo nuclei treated with DMSO (green circles) or 25 µM triptolide (green triangles). *n* = 3 independent extracts; >86 spindles analyzed per extract. P = 0.0376, 0.0313, and 0.9870, one-way ANOVA followed by Tukey’s multiple-comparisons test. (C) Representative immunoblot of TOPBP1 and Ran in *X. tropicalis* extract following mock depletion with rabbit IgG, TOPBP1 immunodepletion, or addback of recombinant TOPBP1 following TOPBP1 immunodepletion. (D) Representative images of metaphase spindles following mock depletion, TOPBP1 immunodepletion, or recombinant TOPBP1 addback following TOPBP1 immunodepletion in species-matched (blue) or hybrid (orange) extract reactions. γH2AX, magenta; DNA, blue; tubulin, green. Scale bar, 10 µm. (E) Percentage of metaphase spindle area occupied by DNA in reactions using stage 8 *X. tropicalis* or *X. laevis* embryo nuclei following mock depletion (circles), TOPBP1 immunodepletion (squares), or recombinant *X. laevis* TOPBP1 addback following TOPBP1 immunodepletion (diamonds). Each point represents an individual spindle. *n* = 4 independent extracts; >179 spindles analyzed per extract. P = 0.5340, P < 0.0001, P < 0.0001, and P > 0.9999, one-way ANOVA followed by Tukey’s multiple-comparisons test. (F) γH2AX fluorescence intensity at the metaphase plate in reactions using stage 8 *X. tropicalis* embryo nuclei (blue) or stage 8 *X. laevis* embryo nuclei (green) following mock depletion (circles), TOPBP1 immunodepletion (squares), or recombinant *X. laevis* TOPBP1 addback following TOPBP1 immunodepletion (diamonds). Each point represents an individual spindle. *n* = 3 independent extracts; >226 spindles analyzed per extract. P = 0.0017, P < 0.0001, P = 0.0013, P < 0.0001, and P < 0.0001, one-way ANOVA followed by Tukey’s multiple-comparisons test. Line indicates the mean, and the error bars indicate one standard deviation above and below the mean.

### TOPBP1 promotes chromosome alignment and DNA damage processing in hybrid reactions

To determine the functional significance of TOPBP1 recruitment in hybrid reactions, we immunodepleted the protein from *X. tropicalis* extract using a *Xenopus*-specific antibody, reducing its levels by >95% (Figure 4C) (Terui et al., 2024). TOPBP1 depletion had little effect on spindle assembly or chromosome alignment in species-matched reactions (Figure 4D,E). In contrast, TOPBP1-depleted hybrid reactions exhibited severe chromosome alignment defects, with DNA distributed throughout the spindle rather than aligned at the metaphase plate (Figure 4D,E). Addition of recombinant *Xenopus* TOPBP1 restored chromosome alignment, demonstrating that TOPBP1 promotes the alignment of damaged, acentric chromosomes in hybrid extract reactions.

We also tested whether TOPBP1 contributes to DNA damage processing in hybrid reactions. TOPBP1 depletion caused a modest increase in γH2AX in species-matched reactions but a substantially greater increase in hybrid reactions (Figure 4D,F). Adding back recombinant TOPBP1 to endogenous levels partially reduced the elevated γH2AX staining, suggesting that TOPBP1 promotes DNA damage processing, although this function was not fully rescued by the recombinant protein. Because RNAPII inhibition reduced TOPBP1 recruitment (Figure 3), we next asked whether TOPBP1 functions upstream or downstream of transcription-associated chromosome damage. Triptolide treatment of TOPBP1-depleted hybrid reactions restored chromosome alignment and significantly reduced γH2AX staining (Figure S4A–C). These results place TOPBP1 downstream of RNAPII activity in the response to chromosome damage in hybrid reactions.

### TOPBP1 and POLθ-dependent DNA repair promote CENP-A removal

We next asked whether TOPBP1-dependent damage responses influence CENP-A loss. Chromosomes isolated from mock-depleted, TOPBP1-depleted, and TOPBP1 addback reactions were analyzed for CENP-A retention. TOPBP1 depletion had no effect on CENP-A in species-matched reactions. Strikingly, however, depletion of TOPBP1 prevented CENP-A loss from *X. laevis* chromosomes exposed to *X. tropicalis* metaphase extract (Figure 5A,B). Addition of recombinant TOPBP1 after depletion failed to restore CENP-A loss, indicating that TOPBP1 is necessary but that the addition of recombinant TOPBP1 to depleted extracts is insufficient to reconstitute this process. Thus, although TOPBP1 depletion increased γH2AX fluorescence intensity (Figure 4F), it prevented CENP-A loss, indicating that DNA damage alone is insufficient to destabilize the centromere and suggesting that TOPBP1-dependent processing of this damage promotes CENP-A removal.

**Figure 5.**
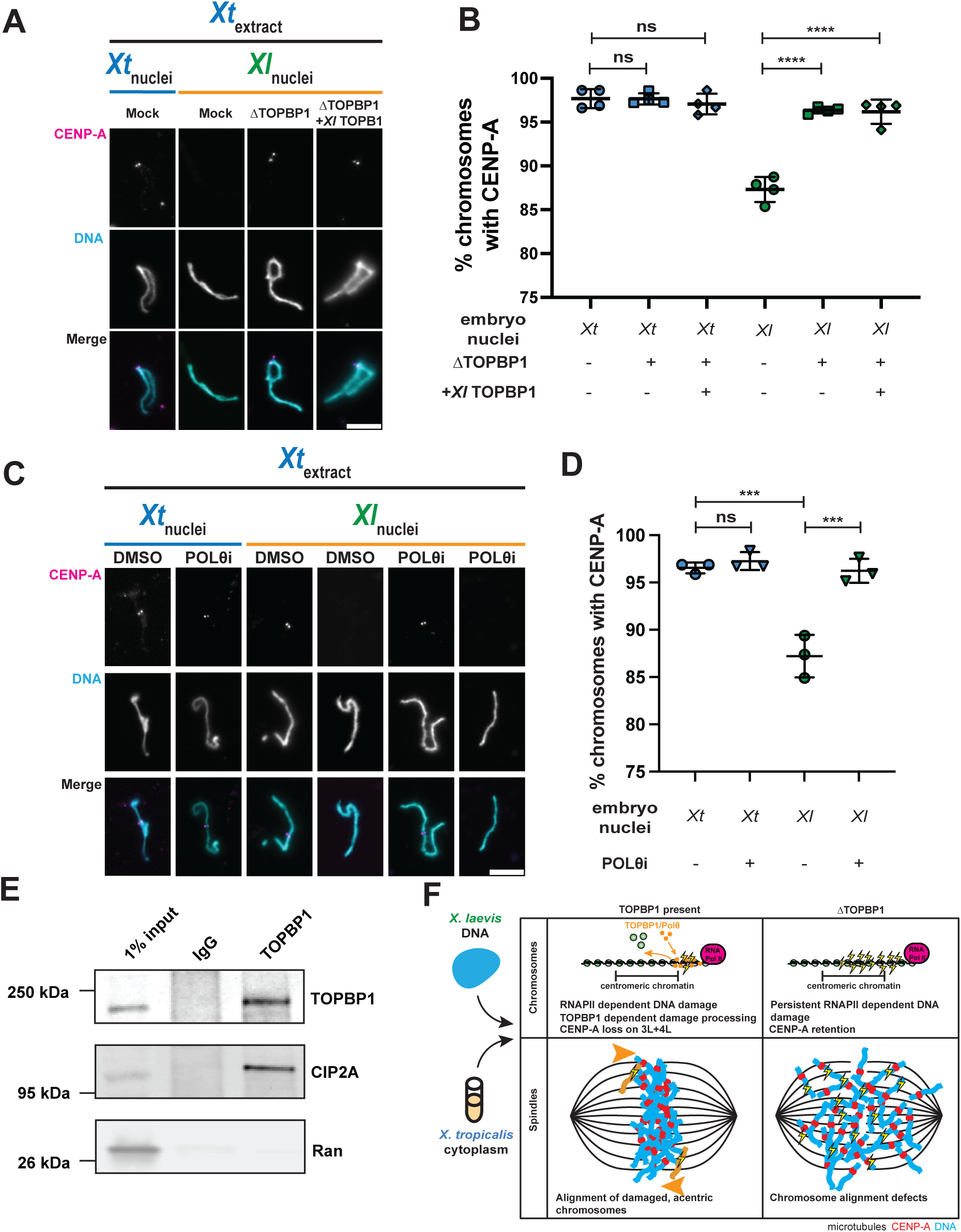
TOPBP1 promotes POLθ-dependent CENP-A removal. (A) Representative images of mitotic chromosomes from stage 8 embryo nuclei following mock depletion, TOPBP1 immunodepletion, or recombinant TOPBP1 addback following TOPBP1 immunodepletion in species-matched (blue) or hybrid (orange) extract reactions. CENP-A, magenta; DNA, cyan. Scale bar, 10 µm. (B) Percentage of mitotic chromosomes positive for CENP-A in reactions using stage 8 *X. tropicalis* embryo nuclei (blue) or stage 8 *X. laevis* embryo nuclei (green) following mock depletion (circles), TOPBP1 immunodepletion (squares), or recombinant *X. laevis* TOPBP1 addback following TOPBP1 immunodepletion (diamonds). *n* = 4 independent extracts; >274 chromosomes analyzed per extract. P > 0.9999, P = 0.9622, P < 0.0001, and P < 0.0001, one-way ANOVA followed by Tukey’s multiple-comparisons test. (C) Representative images of mitotic chromosomes from stage 8 *X. tropicalis* (blue) or *X. laevis* (orange) embryo nuclei cycled into metaphase in *X. tropicalis* extract treated with DMSO or 10 µM ART-558. CENP-A, magenta; DNA, cyan. Scale bar, 5 µm. (D) Percentage of mitotic chromosomes positive for CENP-A in reactions using stage 8 *X. tropicalis* embryo nuclei (blue) or stage 8 *X. laevis* embryo nuclei (green) treated with DMSO (circles) or 10 µM ART-558 (inverted triangles). *n* = 3 independent extracts; >240 chromosomes analyzed per extract. P = 0.9200, P = 0.0002, and P = 0.0002, one-way ANOVA followed by Tukey’s multiple-comparisons test. Line indicates the mean, and the error bars indicate one standard deviation above and below the mean. (E) Representative immunoblot of TOPBP1, CIP2A, and Ran in 1% input, mock immunoprecipitation with rabbit IgG, or TOPBP1 immunoprecipitation. (F) Model illustrating the effects of *X. laevis* chromosomes on spindle assembly and CENP-A retention in *X. tropicalis* cytoplasm in the presence (left) or absence (right) of TOPBP1.

Because hybrid extract reactions are analyzed in metaphase, DNA lesions present at this cell cycle stage must be processed by pathways that remain active when homologous recombination and classical non-homologous end joining are suppressed (Gelot et al., 2023). Recent studies demonstrate that TOPBP1 promotes mitotic DNA repair by recruiting DNA polymerase theta (POLθ) to lesions, facilitating MMEJ and other repair processes (Gelot et al., 2023; Martin et al., 2025). We therefore tested whether POLθ activity contributes to DNA damage processing and CENP-A loss in hybrid extract reactions. Inhibition of POLθ polymerase activity with ART-558 did not alter chromosome morphology or CENP-A stability in species-matched reactions but significantly increased CENP-A retention on *X. laevis* embryo chromosomes in hybrid reactions compared with vehicle-treated controls (Figure 5C,D). These results implicate POLθ-dependent DNA repair in promoting CENP-A loss, consistent with processing of DNA lesions that persist into metaphase in hybrid extract reactions.

Finally, we asked whether POLθ-dependent repair contributes to the chromosome alignment function of TOPBP1. POLθ inhibition increased γH2AX levels in both species-matched and hybrid reactions, although the increase was less pronounced than after TOPBP1 depletion (Figure S5A,B; Figure 4D,F). Despite elevated DNA damage, POLθ inhibition did not impair chromosome alignment (Figure S5A,C). Thus, POLθ-dependent repair contributes to DNA damage processing and CENP-A loss but is dispensable for TOPBP1-mediated chromosome alignment, distinguishing these two functions of TOPBP1.

Consistent with a broader role for TOPBP1 in DNA damage processing, we identified cellular inhibitor of PP2A (CIP2A), a factor implicated in tethering damaged chromosome fragments during mitosis (Adam et al., 2021; De Marco Zompit et al., 2022; Lin et al., 2023; Trivedi et al., 2023), as a TOPBP1-associated protein by immunoprecipitation followed by mass spectrometry (IP-MS). CIP2A was specifically recovered with TOPBP1 relative to IgG controls, and this interaction was validated by Western blotting of TOPBP1 immunoprecipitates (Figure 5E, Table S1). Recent work has implicated TOPBP1 and CIP2A in clustering mitotic chromosome fragments and promoting mitotic DNA repair through pathways including mitotic DNA synthesis (MiDAS) and MMEJ (Martin et al., 2025). This association further supports a role for TOPBP1 in mitotic DNA damage processing and provides a potential explanation for why recombinant TOPBP1 restored chromosome alignment but not γH2AX levels or CENP-A loss after immunodepletion, if CIP2A or other associated repair factors were co-depleted.

Together, these findings identify TOPBP1 as a central mediator of the response to *X. laevis* chromosomes challenged with *X. tropicalis* extract. TOPBP1 promotes alignment of damaged chromosomes while also facilitating their POLθ-dependent damage processing and CENP-A loss. Because TOPBP1 depletion and POLθ inhibition prevent CENP-A loss despite increased DNA damage, DNA damage alone is insufficient to trigger centromere destabilization. Instead, these results support separable roles for TOPBP1 in hybrid chromosome surveillance: promoting the alignment of damaged, acentric chromosomes and facilitating POLθ-dependent DNA damage processing associated with CENP-A removal (Figure 5F).

## Discussion

Hybrid incompatibility is frequently associated with chromosome instability and embryonic lethality, yet how species-specific chromosome defects engage endogenous genome surveillance pathways in the early embryo has remained unclear. Here, we show that centromere instability in *Xenopus* hybrids does not arise from defective CENP-A assembly, but instead results from an active process requiring RNAPII activity and DNA damage processing. We identify TOPBP1 as a central mediator, functioning downstream of transcription-dependent damage to promote acentric chromosome alignment and CENP-A removal through POLθ-dependent DNA repair. Thus, hybrid chromosomes engage a conserved genome surveillance pathway that normally organizes damaged chromosomes and coordinates DNA repair, but in this context, promotes centromere destabilization and chromosome instability.

Our findings revise the current model for centromere defects in *Xenopus* hybrids. Previous studies showed that CENP-A is selectively lost from paternal chromosomes 3L and 4L in hybrid embryos and extract reactions and that supplementation with CENP-A and HJURP rescues this phenotype (Gibeaux et al., 2018; Kitaoka et al., 2022), leading to the proposal that hybrid incompatibility results from defective CENP-A assembly and/or maintenance. By separating interphase CENP-A loading from subsequent metaphase events, we demonstrate that preassembled CENP-A is actively removed after chromosomes encounter *X. tropicalis* cytoplasm. This contrasts with findings in other organisms, where co-evolution of centromere assembly components generates incompatibilities between closely related species, and misregulation of centromeric cohesion contributes to F1 hybrid female sterility in mice (Dudka et al., 2025; El Yakoubi et al., 2026; Maheshwari et al., 2015; Rosin and Mellone, 2016). Together, these studies highlight epigenetic centromere maintenance as a recurrent challenge to successful hybridization.

Our work also reveals an unexpected relationship between DNA damage processing and centromere maintenance. Centromeres experience replication stress and require specialized repair mechanisms to preserve chromosome integrity (Graham et al., 2025; Saayman et al., 2023), but DNA damage has generally been viewed as a consequence of centromere dysfunction with CENP-A loss impairing centromere replication and promoting chromosome rearrangements (Giunta and Funabiki, 2017; Giunta et al., 2021). Here, we show that CENP-A removal requires RNAPII and depends on TOPBP1 and POLθ-dependent DNA damage processing. Importantly, damage alone is insufficient, as TOPBP1 depletion increased γH2AX while preventing centromere destabilization and POLθ inhibition similarly increased damage while promoting CENP-A retention. Thus, the cellular response to DNA damage, rather than damage itself, promotes CENP-A removal, consistent with recent work showing that prolonged replication stress can trigger ATR-dependent CENP-A eviction (Ostapenko et al., 2025). We propose that RNAPII-dependent damage engages TOPBP1-dependent repair machinery and that repair-associated chromatin remodeling or DNA synthesis may destabilize CENP-A-containing chromatin. Although the molecular basis of CENP-A removal remains unknown, these findings suggest that DNA damage processing can itself become a source of centromere instability.

What initiates this pathway selectively on chromosomes 3L and 4L remains an important question. Our data suggest that chromosome specificity is established upstream of TOPBP1 recruitment, as RNAPII inhibition prevents both TOPBP1 localization and CENP-A removal. Moreover, TOPBP1 depletion caused widespread DNA damage while preventing CENP-A loss, indicating that DNA damage alone does not determine which centromeres become destabilized. Chromosome-specific features may therefore render the centromeres of 3L and 4L particularly sensitive to transcription-dependent damage and its processing. Centromeric transcription must be tightly regulated to preserve CENP-A chromatin (Bouzinba-Segard et al., 2006; Chabot et al., 2024; Chan et al., 2012; Grenfell et al., 2016; McNulty et al., 2017), raising the possibility that centromeric or pericentromeric sequences of 3L and 4L are uniquely susceptible to aberrant transcription in *X. tropicalis* cytoplasm. Species-specific differences in repetitive DNA organization, chromatin composition (Long et al., 2023), transcriptional regulation, or replication dynamics could selectively generate or alter the processing of lesions near these centromeres, thereby engaging TOPBP1-dependent pathways that promote CENP-A loss. This resembles inviable *Drosophila* hybrids, in which divergence of a 359-bp satellite block on the paternal X chromosome causes chromosome-specific segregation failure (Ferree and Barbash, 2009). Thus, divergence of repetitive or centromere-associated sequences may render specific chromosomes susceptible to CENP-A loss and centromere destabilization.

Our findings also provide a framework for understanding the fate of *X. laevis* chromosomes 3L and 4L and the low frequency of micronuclei in inviable hybrids during embryogenesis. Previous genome sequencing revealed selective loss of their long arms, while centromeres and short arms are maintained (Gibeaux et al., 2018). Our results suggest that CENP-A removal does not immediately cause chromosome missegregation. Instead, TOPBP1 promotes alignment and clustering of damaged acentric chromosomes, potentially maintaining their ability to segregate. Its chromosome-wide localization on acentrics differs from TOPBP1’s punctate or filamentous localization at mitotic DNA lesions and ultrafine bridges (Bagge et al., 2025; Broderick et al., 2015; Leimbacher et al., 2019; Pedersen et al., 2015) and more closely resembles TOPBP1 localization on acentric chromosomes contained in micronuclei upon mitotic entry in human cells (Lin et al., 2023; Trivedi et al., 2023). TOPBP1 may therefore perform two separable functions: TOPBP1- and POLθ-dependent processing may contribute to CENP-A loss and chromosome rearrangements, whereas TOPBP1 independently promotes alignment of damaged, acentric chromosomes. Its scaffolding and self-oligomerizing activities, together with regulation by PLK1 and CDK1, could facilitate chromosome-wide TOPBP1 localization and the association of acentric chromosomes with the spindle (Balbo Pogliano et al., 2022; Kim et al., 2020; Li et al., 2026). Analogous mechanisms operate in *Drosophila*, where DNA tethers containing BubR1, Polo kinase, INCENP, and Aurora-B connect acentric and centric chromatin fragments and promote their segregation (Blackford and Stucki, 2020; Royou et al., 2010).

More broadly, our findings suggest that reproductive isolation can arise not only from divergence between parental genomes but also from how chromosomes are recognized and processed in a divergent cytoplasmic environment. TOPBP1 and POLθ normally preserve chromosome integrity by organizing damaged chromosomes and promoting DNA repair. In the hybrid context, however, engagement of these conserved pathways can instead promote CENP-A loss and chromosome-specific instability.

## Materials and Methods

### *Xenopus* husbandry

Mature *X. tropicalis* and *X. laevis* were obtained from *Xenopus*1 or the National *Xenopus* Resource (Woods Hole) and maintained and used according to protocols approved by the UC Berkeley Animal Care and Use Committee.

### Egg extract preparation

*X. laevis* cytoplasmic CSF egg extracts were prepared as previously described (Hannak and Heald, 2006; Maresca and Heald, 2006). Briefly, metaphase II-arrested eggs were packed using a tabletop centrifuge prior to centrifugation for 15 minutes at 10,200 rpm at 16°C in a Sorvall HB-6 rotor. Cytoplasm was isolated and supplemented with 10 µg/mL leupeptin, pepstatin, and chymostatin (LPC), 20 µg/mL cytochalasin B, and 1X energy mix (3.75 mM creatine phosphate, 0.5 mM ATP, 0.05 mM EGTA, and 0.5 mM MgCl_2_) and subsequently stored on ice. To visualize microtubules, rhodamine-labeled tubulin was added to reactions at a concentration of 0.3 µM. For immunodepletion experiments, rhodamine-labeled tubulin was added after immunodepletion was complete and a small portion of extract had been reserved for Western blot analysis.

Modifications for *X. tropicalis* cytoplasmic extract included: injecting *X. tropicalis* females with 10 units of human chorionic gonadotropin hormone (hCG) 16-18 hours before use, an additional injection of 250 units of hCG the morning of the experiment, collecting eggs by gentle squeezing, doubling the concentrations of EGTA (10 mM) and MgCl_2_ (2 mM), and storing extract at 16°C following collection (Brown et al., 2007).

### In vitro fertilizations and preparation of Stage 8 embryo nuclei

*X. laevis* stage 8 embryos were generated as previously described (Zhou et al., 2023). In vitro fertilizations to obtain *X. tropicalis* stage 8 embryo nuclei included the following modifications: *X. tropicalis* females were injected with 10 units of hCG 16-18 hours before use and additionally injected with 250 units of hCG the morning of fertilization experiments. *X. tropicalis* males were injected with 250 units of hCG 12-24 hours before dissection. Once females began laying eggs, males were dissected. Testes were collected and stored in Leibovitz L-15 Medium containing 10% fetal bovine serum for use the day of fertilizations.

For all fertilizations, eggs from female frogs were obtained by gentle squeezing onto a petri dish. Testes were homogenized using scissors and a plastic pestle. Eggs were fertilized with 500 µL of sperm solution per dish for 5 minutes. Dishes were flooded with ddH_2_O and incubated for 10 minutes.1/10X MMR was exchanged for ddH_2_O and incubated for 10 minutes. Eggs were incubated with 3% cysteine to remove jelly coats for no longer than 10 minutes. Eggs were then washed in 1/10X MMR and incubated at 23°C for 1-1.5 hours post fertilization. Fertilized embryos that had successfully completed the first cleavage division were sorted for subsequent development preceding isolation of embryo nuclei.

To isolate embryo nuclei, embryos were arrested in interphase by exchange into 1/10X MMR supplemented with 300 µg/mL cycloheximide once they had reached the desired developmental stage. After one hour in cycloheximide solution, embryos were exchanged into Egg Lysis Buffer (ELB) (250 mM sucrose, 50 mM KCl, 2.5 mM MgCl2, 10 mM HEPES pH 7.8) containing 10 µg/mL LPC and 20 µg/mL cytochalasin B, transferred to a 1.5 mL Eppendorf, packed by micro centrifuging tubes on a tabletop centrifuge at 200 g for 1 minute, crushed with a pestle and centrifuged at 10,000 g for 10 minutes at 16°C. Cytoplasmic extract containing nuclei was isolated and gently fixed. Nuclei concentrations were estimated by counting in a Hoechst-stained 0.5 µL drop. Glycerol was added to 8% of the final volume, nuclei were aliquoted, flash frozen in liquid nitrogen, and stored at -80°C.

### Formation of spindles, chromosomes, and nuclei in extract

Replicated sperm nuclei were formed as previously described (Zhou et al., 2023). Briefly, sperm nuclei were added to 25-50 µL of either *X. laevis* or *X.tropicalis* egg extract treated with calcium solution to induce interphase (Hannak and Heald, 2006; Maresca and Heald, 2006). Completion of interphase was determined once nuclei had become swollen 45-70 minutes after initiating interphase reactions. Interphase reactions were driven into metaphase by the addition of fresh, metaphase arrested *X. laevis* or *X. tropicalis* extract. Reactions were incubated at room temperature for 45-60 minutes or until bipolar spindles had formed and chromosomes had condensed. Spindles and chromosomes were fixed and spun down onto coverslips for immunofluorescence as described below.

To examine embryo nuclei in extract reactions, interphase-arrested stage 8 embryo nuclei were thawed on ice for at least 10 minutes, resuspended in 1 mL of CSF-XB containing 10 µg/mL LPC, pelleted at 1600 g for 5 min at 16°C, and gently resuspended in a minimal volume of CSF-XB containing 10 µg/mL LPC. To examine nuclei in interphase, nuclei were incubated in *X. tropicalis* extract arrested in interphase via the addition of calcium solution (Hannak and Heald, 2006; Maresca and Heald, 2006) for 15 minutes prior to being fixed and spun down onto coverslips for subsequent immunofluorescence as described below. To examine metaphase spindles and chromosomes generated from embryo nuclei, nuclei were incubated with 25-50 µL of metaphase-arrested *X. tropicalis* extract at room temperature for 45-60 minutes or until bipolar spindles had formed and chromosomes had condensed. Spindles and chromosomes were fixed and spun down onto coverslips for subsequent immunofluorescence as described below.

### Chromosome spin-downs, immunofluorescence, and quantification

Metaphase chromosomes from spindle reactions were spun down and processed for immunofluorescence as described previously (Brown et al., 2007; Maresca and Heald, 2006). Briefly, extract reactions were diluted into 200 µL of chromosome dilution buffer (250 mM sucrose, 10 mM HEPES pH 8.0, 0.5 mM EGTA, 200 mM KCl, 1 mM MgCl_2_) for 10 minutes, then fixed in 5 mL of chromosome fixation buffer (5 mM HEPES pH 7.8, 0.1 mM EDTA, 100 mM NaCl, 2 mM KCl, 1 mM MgCl_2_, 2 mM CaCl_2_, 0.5% Triton X-100, 20% glycerol, and 2% paraformaldehyde) for 10 minutes. Fixed reactions were then layered on a 5 mL cushion (5 mM HEPES pH 7.8, 0.1 mM EDTA, 100 mM NaCl, 2 mM KCl, 1 mM MgCl_2_, 2 mM CaCl_2_, 40% glycerol) and spun down at 5,500 rpm using a Sorvall HS-4 rotor for 20 minutes at 16°C. Coverslips were removed and additionally fixed for 30 seconds in ice-cold methanol, washed three times in PBS + 0.1% NP40, and blocked overnight in PBS + 3% BSA. Coverslips were then quickly washed with PBS + 0.1% NP40 once prior to incubation with primary antibodies diluted in PBS + 3% BSA for 1 hour. Primary antibodies were used at the following concentrations: rabbit anti-xCENP-A (1:500; Straight lab; Stanford), rabbit anti-xTOPBP1 (1:1000; Chistol lab; Stanford), rabbit anti-xCENP-T-488 (1:500; Straight lab; Stanford). After washing with PBS + 0.1% NP40 coverslips were incubated with 1:1000 anti-rabbit secondary antibodies coupled to Alexa Fluor 488 or 647 for 30 minutes and then with 1:1000 Hoechst for 5 minutes. Coverslips were washed with PBS + 0.1% NP40 and mounted using ProLong Gold. For experiments utilizing rabbit anti-xTOPBP1 and rabbit anti-xCENP-T directly conjugated to Alexa-fluor 488, coverslips were first incubated with rabbit anti-xTOPBP1, washed, incubated with goat anti-rabbit Alexa fluor 647 secondary antibody, washed, and blocked with PBS-rabbit IgG for one hour, and washed with PBS + 0.1% NP40 prior to incubation with rabbit anti-xCENPT-488. Quantification of CENP-A, CENP-T, and TOPBP1 localization was determined manually using FIJI for each experiment. Only single chromosomes were counted. The average of each extract was calculated in Microsoft Excel as a percentage of total chromosome number for a given condition. Averages for each experiment were plotted using Prism 11.0.

### Spindle spin-downs, immunofluorescence, and quantification

Metaphase spindles from spindle reactions were spun down and processed for immunofluorescence as described previously (Brown et al., 2007; Hannak and Heald, 2006; Maresca and Heald, 2006). Briefly, extract reactions were fixed in 1 mL of spindle dilution buffer (30% glycerol in BRB80 containing 2.5% paraformaldehyde) for 10 minutes. Fixed reactions were layered on top of 5 mL of spindle cushion (40% glycerol in BRB80) and spun down onto coverslips at 5,500 rpm using a Sorvall HS-4 rotor for 20 minutes at 16°C. Coverslips were removed and additionally fixed for 5 minutes in ice-cold methanol, washed three times in PBS + 0.1% NP40, and blocked overnight in PBS + 3% BSA. Coverslips were then quickly washed with PBS + 0.1% NP40 once prior to incubation with primary antibodies diluted in PBS + 3% BSA for one hour. Antibodies were used at the following concentrations: rabbit anti-xCENPA (1:500; Straight lab; Stanford), rabbit anti-TOPBP1 (1:500; AB105109; Abcam), rabbit anti-xCENPT-488 (1:500; Straight lab; Stanford), mouse anti-phospho-Histone H2A.X (Ser139), clone JBW301 (1:250; 05-636-25UG; MilliporeSigma). After washing with PBS + 0.1% NP40 coverslips were incubated with 1:1000 anti-rabbit secondary antibodies coupled to Alexa Fluor 488 or 647 for 30 minutes and then with 1:1000 Hoechst for 5 minutes. Coverslips were washed with PBS + 0.1% NP40 and mounted using ProLong Gold. For experiments utilizing rabbit anti-xTOPBP1 and rabbit anti-xCENPT directly conjugated to Alexa-fluor 488, coverslips were first incubated with rabbit anti-xTOPBP1, washed, incubated with goat anti-rabbit Alexa fluor 647 secondary antibody, washed, blocked with PBS-rabbit IgG for one hour, and washed prior to incubation with rabbit anti-xCENPT-488.

Quantification of TOPBP1, γH2AX, and CENP-T localization to chromosomes in metaphase spindles was done manually in FIJI by examining sum projections of metaphase spindles from extract reactions. The average of each extract was calculated as a percentage of the total number of spindles for a given experimental condition. Only single spindles were quantified.

To quantify fluorescence intensity of TOPBP1 at spindle poles, total fluorescence intensity of TOPBP1 from sum projections generated in FIJI was measured in a square (3.7 × 3.7 microns) centered on spindle poles. To quantify fluorescence intensity of TOPBP1 throughout metaphase spindles, total fluorescence intensity of TOPBP1 from sum projections generated in FIJI was measured by using the freehand selection tool to trace the rhodamine-tubulin signal for a given spindle to specify a region of interest (ROI) marking the entire spindle. To quantify fluorescence intensity of TOPBP1 and γH2AX on the metaphase plate, total fluorescence intensity of TOPBP1 or γH2AX from sum projections generated in FIJI was measured by using the polygon selection tool to trace the DNA signal in metaphase spindles and specify an ROI encompassing the metaphase plate. Background subtractions from ROIs of the same size and shape were subtracted from these measurements for each type of quantification in each image. Only single spindles were quantified.

To quantify the percentage of a given spindle occupied by DNA, sum projections of metaphase spindles were generated in FIJI. The polygon selection tool was used to trace the DNA signal in each spindle and specify an ROI defining the location of DNA throughout the spindle. The freehand selection tool was used similarly to trace the rhodamine-tubulin signal and specify an ROI defining the spindle. The area of each ROI was measured and the area of the DNA signal was divided by the area of the tubulin signal and multiplied by 100 to determine the relative proportion of the spindle area occupied by DNA

### Nuclei spin-downs, immunofluorescence, and quantification

Interphase nuclei from extract reactions were spun down and processed for immunofluorescence as described previously. Briefly, reactions were fixed in 1 mL of nuclei dilution buffer consisting of ELB supplemented with 15% glycerol and 2.6% paraformaldehyde for 10 minutes. Fixed reactions were layered on top of 5 mL of nuclei cushion buffer (200 mM Sucrose,100 mM KCl,1 mM MgCl2,10 mM HEPES pH 7.8,0.1 mM CaCl2, 25% glycerol) and spun down onto coverslips at 5,500 rpm using a Sorvall HS-4 rotor for 20 minutes at 16°C. Coverslips were removed and additionally fixed for 5 minutes in ice-cold methanol, washed three times in PBS + 0.1% NP40, and blocked overnight in PBS + 3% BSA. Coverslips were then quickly washed with PBS + 0.1% NP40 once prior to incubation with primary antibodies diluted in PBS + 3% BSA for one hour. The following antibodies were used at the following concentrations: rabbit anti-xCENPA (1:500; Straight lab; Stanford). After washing with PBS + 0.1% NP40 coverslips were incubated with 1:1000 anti-rabbit secondary antibodiy coupled to Alexa Fluor 488 for 30min and then with 1:1000 Hoechst for 5 minutes. Coverslips were washed with PBS + 0.1% NP40 and mounted using ProLong Gold. Quantification of CENP-A localization was determined manually using FIJI for each experiment. Sum projections of nuclei were generated in FIJI. The multi-point tool was then used to count the number of CENP-A foci per nucleus. The average number of CENP-A foci for a given experimental condition was calculated and compared to the expected number of CENP-A foci for *X. tropicalis* or *X. laevis* nuclei in Microsoft Excel. Only single nuclei were counted. The average percentage of CENP-A foci per nucleus relative to the expected number of CENP-A foci and the number of CENP-A foci quantified in each nucleus for each experiment was pooled and plotted using Prism 11.0.

### Drug treatments and protein additions

*X. tropicalis* and *X. laevis* extracts were supplemented with the following drugs and concentrations: Triptolide (RNAPII inhibitor, 25 µM), α-amanitin (RNAPII inhibitor, 50 µg/ml), ART-558 (POLθ inhibitor, 10 µM)

For TOPBP1 addback experiments following immunodepletion, recombinant *X. laevis* TOPBP1 (Terui et al., 2024) was added to extracts at a concentration of 35 nM based on estimated abundance from previous mass spectrometry data from *X. laevis* (Wühr et al., 2014). Extracts supplemented with recombinant *X. laevis* TOPBP1 were examined by Western blot to determine whether the concentration of TOPBP1 in supplemented extracts was similar to endogenous levels of TOPBP1 in mock depleted control extracts.

For experiments supplemented with centromere assembly factors, a final volume equivalent to 2% of the egg extract reaction volume of HJURP, CENP-A, or both protein products from TnT Sp6-coupled rabbit reticulocyte in vitro transcription/translation reactions was added to extract reactions prior to the addition of DNA (replicated sperm nuclei or interphase arrested embryo nuclei) (Kitaoka et al., 2022).

### Immunodepletion

Immunodepletion of TOPBP1 from *X. tropicalis* egg extracts was carried out as described previously (Hannak and Heald, 2006) with slight modifications to accommodate differences between *X. tropicalis* and *X. laevis* cytoplasmic extracts. Briefly, 50 µl of Protein A Dynabeads were conjugated to 10 µg of polyclonal rabbit anti-xTOPBP1 antibody or 10 µg rabbit IgG isotype control. 50 µl of extract was incubated with 25 µl of antibody coupled beads for 1 hour at 16°C with gentle mixing every 15 minutes. TOPBP1 or mock depleted extract was isolated from beads using a magnetic bead concentrator prior to spindle assembly extract reactions.

### Immunoprecipitation

12 µg of rabbit anti-xTOPBP1 antibody was coupled to 60 µl of protein A dynabeads as previously described (Hannak and Heald, 2006). For Immunoprecipitation of TOPBP1, 120 µl of *X. tropicalis* extract was subjected to a 1-hour incubation at 16°C with gentle mixing every 15 minutes. Beads were washed extensively with XB before eluting for 5 minutes at 95°C in 1x Laemmli sample buffer and retrieving eluate by separating beads from supernatant on a magnetic bead concentrator.

### Mass Spectrometry

This work was performed by QB3 Mass Spectrometry at the University of California, Berkeley. Immunoprecipitation samples were resolved by SDS-PAGE electrophoresis and stained with Gelcode Blue Coomassie stain (Pierce). Each lane to be analyzed was cut from the gel. Gel slices were cut into 1mm^2^ cubes with a fresh razor blade and transferred to a clean microcentrifuge tube. The gel pieces were washed twice with 50% ACN and 50mM Ammonium Bicarbonate pH 8 for 15 minutes with shaking. The gel pieces were dehydrated with 100% ACN for 5 minutes with shaking. Then the solvent was removed, and the gel pieces were allowed to air dry for 20 min. To dry gel pieces, 10mM TCEP and 40mM CAA were added and incubated at 70°C for 5 minutes. The gel pieces were washed again with 50% ACN and 50% 50mM ABC for 15 minutes with shaking. The gel pieces were rehydrated in 50 mM ABC and 1ug Trypsin/ Lys C (1:50) was added and incubated for 1 hour at room temperature, then 50mM HEPES pH 8 solution was added to cover the pieces. The samples were allowed to incubate overnight at 37°C. Peptides were extracted from gel pieces with 25% ACN and 50mM Ammonium Bicarbonate pH 8 then 100% ACN for 5 minutes with shaking. Samples were filtered through a 0.22µm PVDF spin column (Millipore) Peptides were dried in a speedvac to 30µl and acidified with 2µl formic acid (NEAT).

Trypsin digested peptides were analyzed by online capillary nanoLC-MS/MS using a 25cm reversed phase column and a 10cm precolumn fabricated in-house (75µm inner diameter, packed with ReproSil-Gold C18-1.9μm resin (Dr. Maisch GmbH)) that was equipped with a laser-pulled nanoelectrospray emitter tip. The precolumn used 3.0µm packing (Dr. Maisch GmbH). Peptides were eluted at a flow rate of 300nl/min using a linear gradient of 2–40% buffer B in 140 min (buffer A: 0.05% formic acid and 5% acetonitrile in water; buffer B: 0.05% formic acid and 95% acetonitrile in water) in a Thermo Fisher Easy-nLC1200 nanoLC system. Peptides were ionized using a FLEX ion source (Thermo Fisher) using electrospray ionization into a Fusion Lumos Tribrid Orbitrap Mass Spectrometer (Thermo Fisher Scientific). Data was acquired in orbi-trap mode. Instrument method parameters were as follows: MS1 resolution, 120,000 at 200 m/z; scan range, 350−1600 m/z. The top 20 most-abundant ions were subjected to collision-induced dissociation with a normalized collision energy of 35%, activation q 0.25, and precursor isolation width 2 m/z. Dynamic exclusion was enabled with a repeat count of 1, a repeat duration of 30 seconds, and an exclusion duration of 20 seconds. RAW files were analyzed using PEAKS(Bioinformatics Solution Inc) with the following parameters: semi-specific cleavage specificity at the C-terminal site of R and K, allowing for 4 missed cleavages, precursor mass tolerance of 15 ppm, and fragment ion mass tolerance of 0.5 Daltons. Methionine oxidation (15.99, M), was set as variable modifications and Cysteine carbamidomethylation (57.02, C) was set as a fixed modification. Peptide hits were filtered using a 1% FDR. Label free quantitation (LFQ) was done using PEAKS default settings.

### Western blots

Extract protein concentrations for Western blots testing immunodepletion of TOPBP1 were measured by BCA assay (Thermo Scientific). Samples were prepared in 1x Laemmli sample buffer supplemented with 2.5% beta-mercaptoethanol and loaded to achieve 10 µg/lane. For Western blots of TOPBP1 immunoprecipitates, 1% of the total volume of crude extract used for immunoprecipitation (typically 1.2 µL) was diluted in 1x Laemmli sample buffer to a final volume of 5 µL and compared with an equivalent volume of eluate from IgG or TOPBP1 immunoprecipitations. All samples were boiled at 95°C for 5 minutes, resolved on BioRad Mini PROTEAN TGX 4-20% gradient gels, transferred to nitrocellulose for 2 hours at 60 Volts in transfer buffer with 20% methanol. Membranes were blocked in 5% dry milk, incubated with primary antibodies diluted in 1x PBST + 3% Bovine Serum Albumin overnight at 4°C, washed, and incubated with secondary antibodies diluted in 1x PBST + 4% milk at 1:10,000 for 1 hour at room temperature. The following primary antibodies were used at the following concentrations: rabbit anti-xTOPBP1 (1:5000; Chistol lab; Stanford), rabbit anti-CIP2A(1:250; 23199-1-AP; Proteintech), mouse anti-Ran (1:2000; 610341; BD Transduction Laboratories). The following secondary antibodies were used: Goat anti-Mouse IRDye 680RD (1:10000; 926-68070; LI-COR Biosciences), Goat anti-Rabbit IRDye 800CW (1:10000; 92632211; LI-COR Biosciences). Membranes were imaged on a Li-COR Odyssey CLx. To compare relative amounts of protein between conditions, band intensities were quantified in FIJI. The ratio of TOPBP1 band intensity to Ran band intensity was used to determine whether a given immunodepletion had succeeded.

### Chromosome, spindle, and nuclei imaging

All images were taken using Olympus cellSens Dimension 2 software on an upright Olympus BX53 microscope with an ORCA-Spark camera and Olympus UPlan 60x/NA 1.40 oil objective.

### Statistical tests

Welch’s *t* test and one-way ANOVA, as indicated in figure legends, were performed using Prism 11.0

## Acknowledgements

We thank Maiko Kitaoka, Coral Zhou, Helena Cantwell, Gabriel Cavin, and Alex Lessenger for advice and feedback on the manuscript; Robert Maxwell and Berkeley QB3 for conducting mass spectrometry and analysis (RRID:SCR_025852); Aaron Straight’s lab at Stanford University for providing CENP-A and CENP-T antibodies; We also acknowledge Arielle McMullan for contributing to extract preparation and attempted FISH on mitotic *Xenopus* chromosomes, and current and former Heald Lab members for thoughtful discussions throughout this project This work was supported by the National Institutes of Health MIRA grant R35GM118183 and the Flora Lamson Hewlett Chair in Biochemistry to R. Heald, a Cancer Research Coordinating Committee predoctoral fellowship to C. Clark, an NSF CAREER award (2144481), a NIGMSR35 award (GM147060), and an American Cancer Society seed grant (228425) to G. Chistol. QB3 Mass Spectrometry received support from the National Institutes of Health (grant number 1S10RR025622-01 and 1S10OD020062-01).

## Author contributions

CC: conceptualization, formal analysis, investigation, methodology, visualization, and writing-original draft, review, and editing. RT: resources. GC: resources, writing-review and editing. RH: conceptualization, methodology, funding acquisition, project administration, supervision, writing-original draft, review, and editing.

---

**Figure S1.**
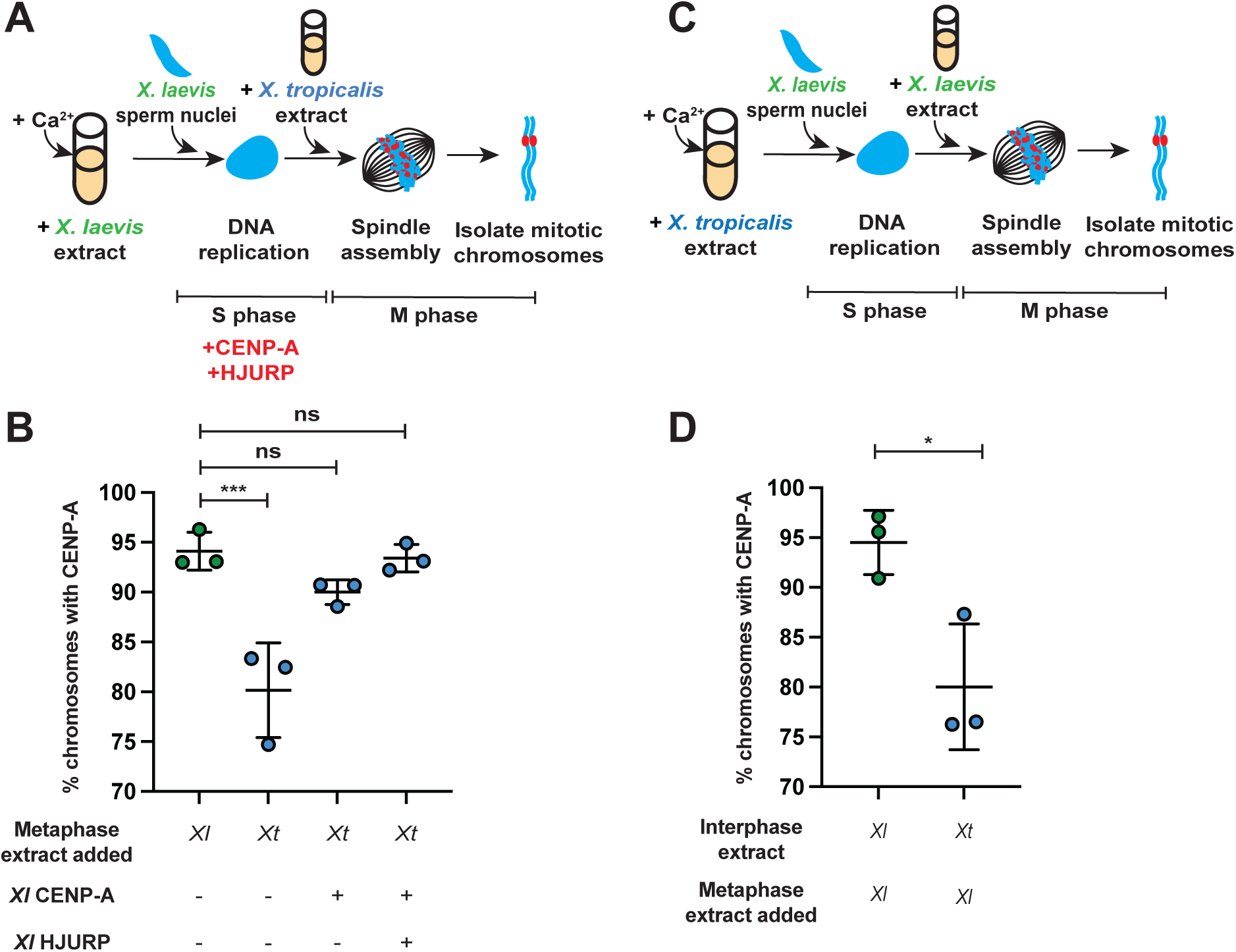
Promoting S-phase centromere assembly stabilizes CENP-A on *X. laevis* chromosomes. (A) Schematic of the hybrid extract reaction supplemented with centromere assembly factors during interphase. *X. laevis* sperm nuclei were replicated in species-matched *X. laevis* interphase extract supplemented with reticulocyte TnT reaction products generated using no DNA control, CENP-A, or CENP-A and HJURP, and subsequently cycled into metaphase in *X. tropicalis* extract. (B) Percentage of replicated *X. laevis* sperm chromosomes positive for CENP-A after cycling into metaphase in *X. tropicalis* extract following interphase in species-matched *X. laevis* extract supplemented with the indicated centromere assembly factors. *n* = 3 independent extracts; >207 chromosomes analyzed per extract. P = 0.0011, 0.3185, and 0.9884, one-way ANOVA followed by Tukey’s multiple-comparisons test. (C) Schematic of the reciprocal hybrid extract reaction. *X. laevis* sperm nuclei were replicated in *X. tropicalis* interphase extract and subsequently cycled into metaphase in *X. laevis* extract. (D) Percentage of replicated *X. laevis* sperm chromosomes positive for CENP-A after cycling into metaphase in the indicated extract reactions. *n* = 3 independent extracts; >133 chromosomes analyzed per extract. P = 0.0387, Welch’s t test. Line indicates the mean, and the error bars indicate one standard deviation above and below the mean.

**Figure S2.**
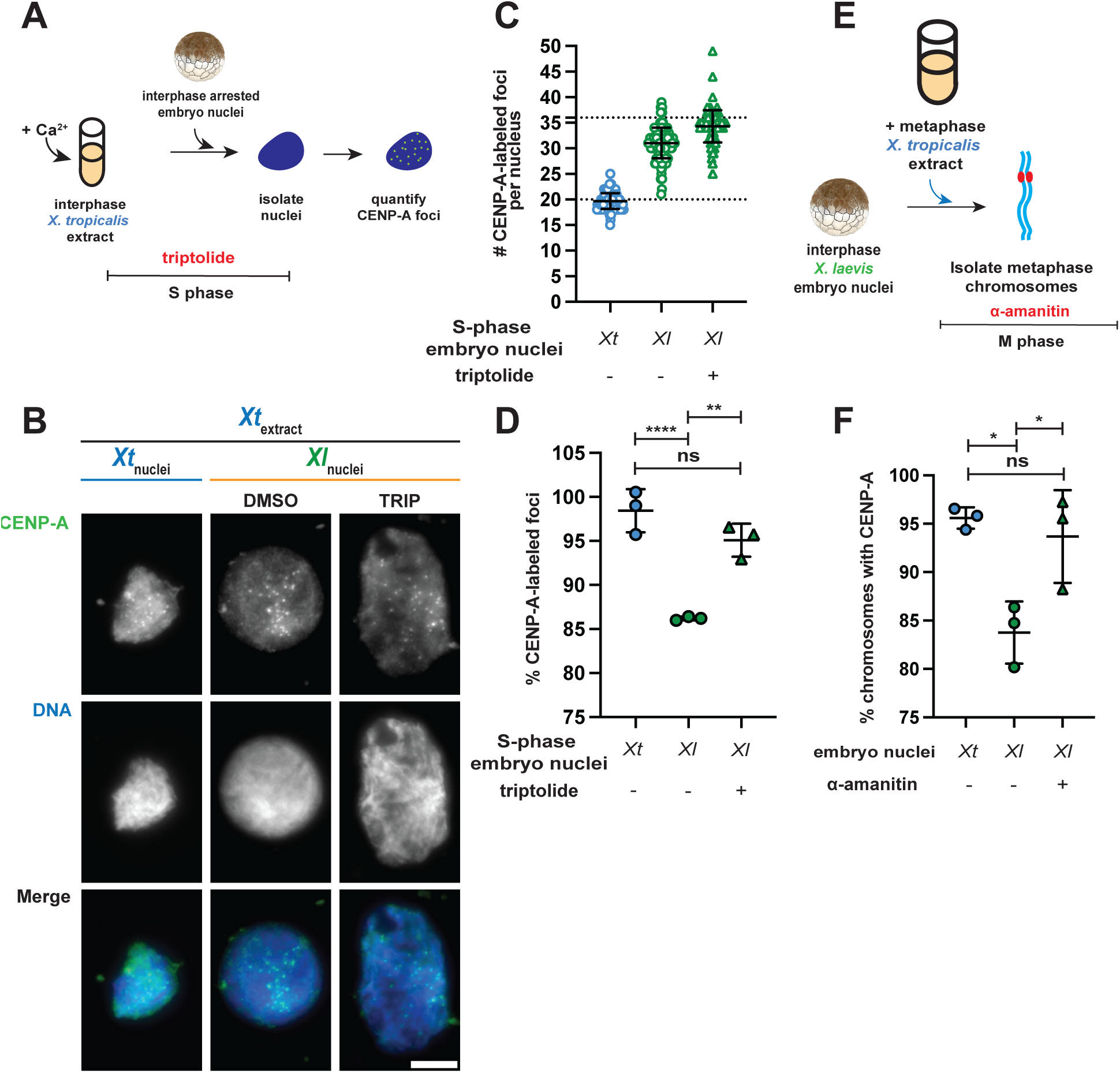
Effects of transcription inhibition on CENP-A retention in hybrid extract reactions. (A) Schematic of interphase extract reactions using stage 8 embryo nuclei treated with triptolide. (B) Representative images of stage 8 *X. tropicalis* or *X. laevis* embryo nuclei incubated in *X. tropicalis* extract treated with DMSO or triptolide. CENP-A (green), DNA (blue). Scale bar, 5 µm. (C) Number of CENP-A–positive foci per nucleus in *X. tropicalis* embryo nuclei incubated in *X. tropicalis* extract (blue circles) or *X. laevis* embryo nuclei incubated in *X. tropicalis* extract treated with DMSO (green circles) or triptolide (green triangles). Each point represents an individual nucleus. Dashed lines at 20 and 36 indicate the expected numbers of CENP-A foci in *X. tropicalis* and *X. laevis* nuclei, respectively. *n* = 3 independent extracts; >116 nuclei analyzed per extract. (D) Percentage of expected CENP-A–positive foci observed in *X. tropicalis* embryo nuclei incubated in *X. tropicalis* extract (blue circles) or *X. laevis* embryo nuclei incubated in *X. tropicalis* extract treated with DMSO (green circles) or triptolide (green triangles). *n* = 3 independent extracts; >116 nuclei analyzed per extract. P = 0.0004, 0.0022, and 0.1344, one-way ANOVA followed by Tukey’s multiple-comparisons test. (E) Schematic of hybrid extract reactions using stage 8 embryo nuclei treated with α-amanitin during metaphase. (F) Percentage of chromosomes positive for CENP-A after *X. tropicalis* embryo nuclei were cycled into metaphase in species-matched *X. tropicalis* extract or *X. laevis* embryo nuclei were cycled into metaphase in *X. tropicalis* extract treated with DMSO or α-amanitin. *n* = 3 independent extracts; >157 chromosomes analyzed per extract. P = 0.0124, 0.7778, and 0.0269, one-way ANOVA followed by Tukey’s multiple-comparisons test. Line indicates the mean, and the error bars indicate one standard deviation above and below the mean.

**Figure S3.**
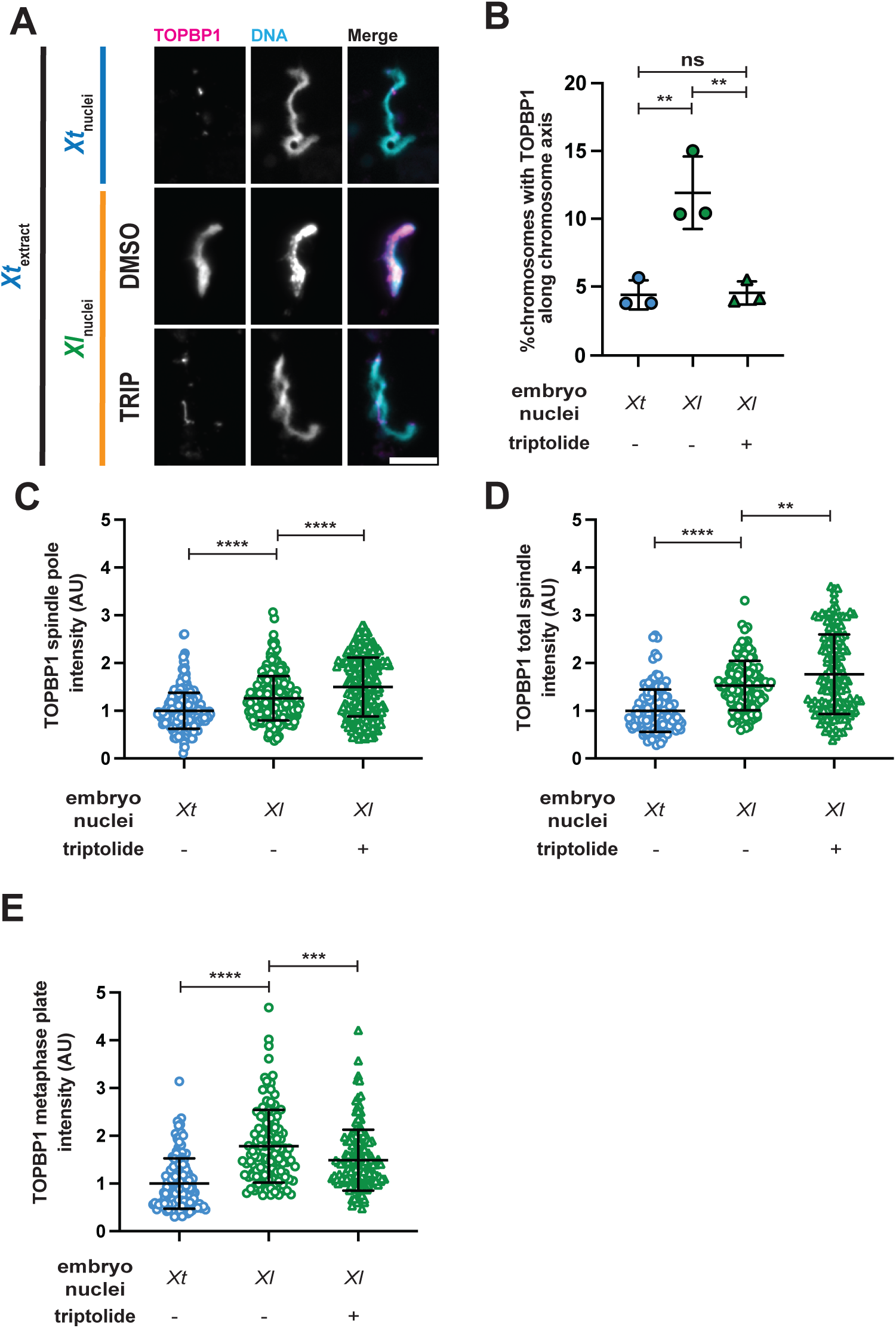
TOPBP1 localization in *X. tropicali*s and hybrid extract reactions. (A) Representative immunofluorescence images of mitotic chromosomes from *X. tropicalis* extract reactions using *X. tropicalis* or *X. laevis* embryo nuclei treated with DMSO or 25 µM triptolide. DNA. TOPBP1 (magenta), DNA (cyan). Scale bar, 5 µm. (B) Percentage of mitotic chromosomes displaying TOPBP1 staining along the chromosome axis in reactions using *X. tropicalis* embryo nuclei (blue circles) or *X. laevis* embryo nuclei treated with DMSO (green circles) or 25 µM triptolide (green triangles). n = 3 independent extracts; >176 chromosomes analyzed per extract. P = 0.0042, 0.0048, and 0.9860, one-way ANOVA followed by Tukey’s multiple-comparisons test. (C) TOPBP1 fluorescence intensity at spindle poles in *X. tropicalis* extract reactions using *X. tropicalis* embryo nuclei (blue circles) or *X. laevis* embryo nuclei treated with DMSO (green circles) or 25 µM triptolide (green triangles). Each point represents an individual spindle pole. n = 3 independent extracts; >274 spindles analyzed per extract. P < 0.0001 and P < 0.0001, one-way ANOVA followed by Tukey’s multiple-comparisons test. (D) TOPBP1 fluorescence intensity across the entire spindle in *X. tropicalis* extract reactions using stage 8 *X. tropicalis* embryo nuclei (blue circles) or stage 8 *X. laevis* embryo nuclei treated with DMSO (green circles) or 25 µM triptolide (green triangles). Each point represents an individual spindle. n = 3 independent extracts; >137 spindles analyzed per extract P < 0.0001 and P = 0.055, one-way ANOVA followed by Tukey’s multiple-comparisons test. (E) TOPBP1 fluorescence intensity at the metaphase plate in *X. tropicalis* extract reactions using stage 8 *X. tropicalis* embryo nuclei (blue circles) or stage 8 *X. laevis* embryo nuclei treated with DMSO (green circles) or 25 µM triptolide (green triangles). Each point represents an individual metaphase plate. n = 3 independent extracts; >137 metaphase plates analyzed per extract. P < 0.0001 and P = 0.0007, one-way ANOVA followed by Tukey’s multiple-comparisons test. Line indicates the mean, and the error bars indicate one standard deviation above and below the mean.

**Figure S4.**
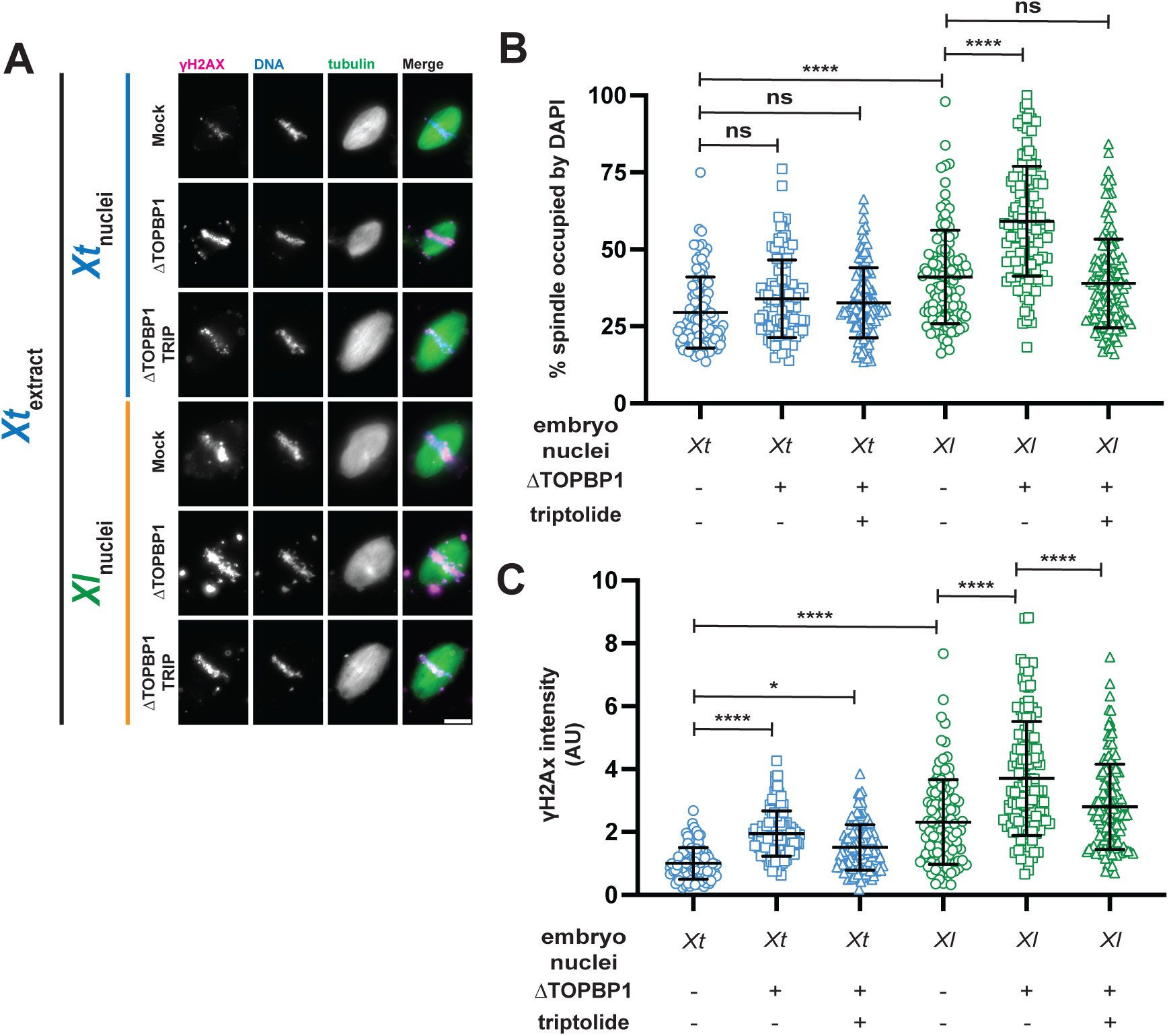
RNA Polymerase II inhibition suppresses chromosome misalignment and DNA damage following TOPBP1 depletion. (A) Representative images of metaphase spindles following mock depletion, TOPBP1 immunodepletion, or treatment with 25 µM triptolide following TOPBP1 immunodepletion in species-matched (blue) or hybrid (orange) extract reactions. γH2AX, magenta; DNA, blue rhodamine-labeled tubulin, green. Scale bar, 10 µm. (B) Percentage of metaphase spindle area occupied by DNA in reactions using stage 8 *X. tropicalis* embryo nuclei (blue) or stage 8 *X. laevis* embryo nuclei (green) following mock depletion (circles), TOPBP1 immunodepletion (squares), or treatment with 25 µM triptolide following TOPBP1 immunodepletion (triangles). Each point represents an individual spindle. *n* = 3 independent extracts; >213 spindles analyzed per extract. P = 0.2117, P = 0.5633, P < 0.0001, P < 0.0001, and P = 0.7123, one-way ANOVA followed by Tukey’s multiple-comparisons test. (C) γH2AX fluorescence intensity at the metaphase plate in reactions using stage 8 *X. tropicalis* embryo nuclei (blue) or stage 8 *X. laevis* embryo nuclei (green) following mock depletion (circles), TOPBP1 immunodepletion (squares), or treatment with 25 µM triptolide following TOPBP1 immunodepletion (triangles). Each point represents an individual spindle. *n* = 3 independent extracts; >213 spindles analyzed per extract. P < 0.0001, P = 0.0105, P < 0.0001, P < 0.0001, and P < 0.0001, one-way ANOVA followed by Tukey’s multiple-comparisons test. Line indicates the mean, and the error bars indicate one standard deviation above and below the mean.

**Figure S5.**
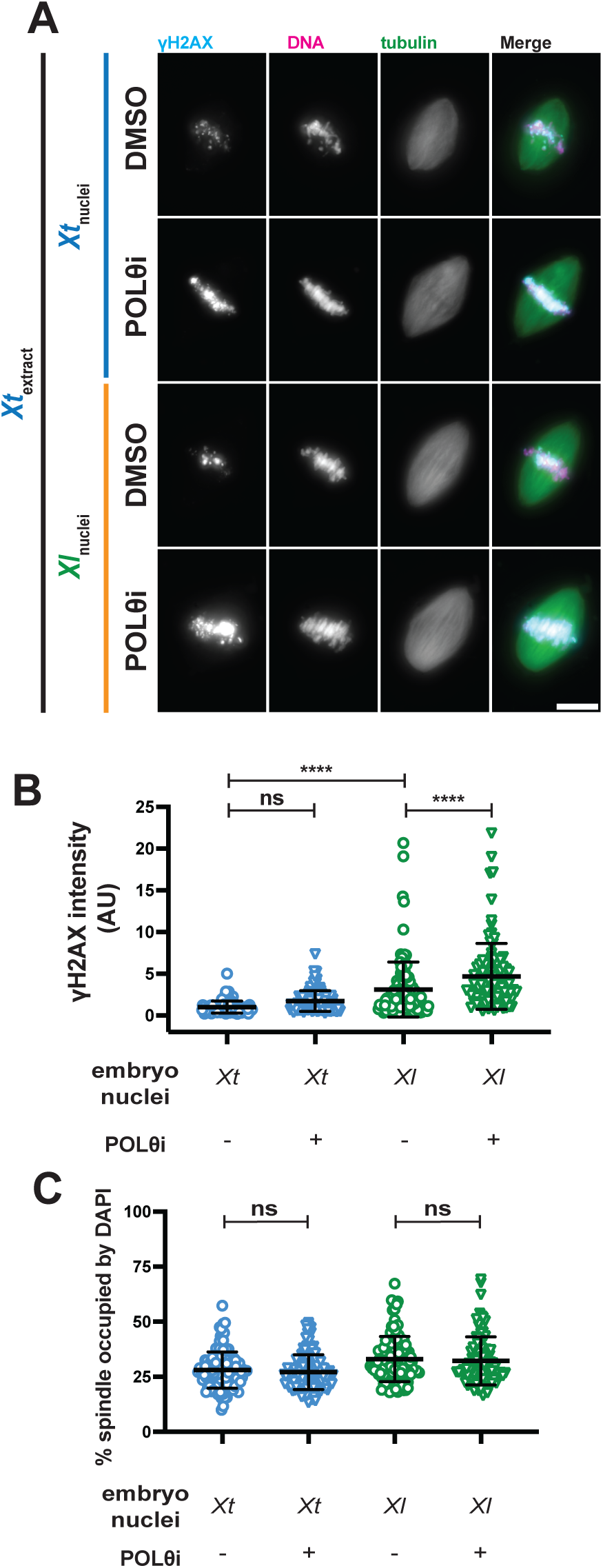
POLθ inhibition exacerbates DNA damage in hybrid extract reactions. (A) Representative images of metaphase spindles assembled in *X. tropicalis* extract using stage 8 *X. tropicalis* embryo nuclei (blue) or stage 8 *X. laevis* embryo nuclei (orange) treated with DMSO or 10 µM ART-558. γH2AX, cyan; DNA, magenta; rhodamine-labeled tubulin, green. Scale bar, 10 µm. (B) Percentage of metaphase spindle area occupied by DNA in reactions using stage 8 *X. tropicalis* embryo nuclei (blue) or stage 8 *X. laevis* embryo nuclei (green) treated with DMSO (circles) or 10 µM ART-558 (inverted triangles). Each point represents an individual spindle. *n* = 3 independent extracts; >129 spindles analyzed per extract. P = 0.8673 and P = 0.9194, one-way ANOVA followed by Tukey’s multiple-comparisons test. (C) γH2AX fluorescence intensity at the metaphase plate in reactions using stage 8 *X. tropicalis* embryo nuclei (blue) or stage 8 *X. laevis* embryo nuclei (green) treated with DMSO (circles) or 10 µM ART-558 (inverted triangles). Each point represents an individual spindle. *n* = 3 independent extracts; >129 spindles analyzed per extract. P = 0.1192, P < 0.0001, and P < 0.0001, one-way ANOVA followed by Tukey’s multiple-comparisons test. Line indicates the mean, and the error bars indicate one standard deviation above and below the mean.

